# Proteins Need Extra Attention: Improving the Predictive Power of Protein Language Models on Mutational Datasets with Hint Tokens

**DOI:** 10.1101/2023.12.05.570055

**Authors:** Xinning Li, Ryann Perez, Sam Giannakoulias, E. James Petersson

## Abstract

In this computational study, we introduce “hint token learning,” a novel machine learning approach designed to enhance protein language modeling. This method effectively addresses the unique challenges of protein mutational datasets, characterized by highly similar inputs that may differ by only a single token. Our research highlights the superiority of hint token learning over traditional fine-tuning methods through three distinct case studies. We first developed a highly accurate free energy of folding model using the largest protein stability dataset to date. Then, we applied hint token learning to predict a biophysical attribute, the brightness of green fluorescent protein mutants. In our third case, hint token learning was utilized to assess the impact of mutations on RecA bioactivity. These diverse applications collectively demonstrate the potential of hint token learning for improving protein language modeling across general and specific mutational datasets. To facilitate broader use, we have integrated our protein language models into the HuggingFace ecosystem for downstream, mutational fine-tuning tasks.

## MAIN

In recent years, large language modeling (LLM)^1-3^ has emerged as a transformative approach in natural language processing (NLP)^4,5^, revolutionizing various tasks such as machine translation^6,7^, text generation^8^, and sentiment analysis.^9^ The introduction of the transformer model by Vaswani *et al*. marked a significant milestone in LLM.^6^ The transformer architecture integrated insights from Recurrent Neural Networks (RNNs)^10^ and Long Short-Term Memory Neural Networks (LSTMs)^11^, while introducing a novel approach, the self-attention mechanism. By harnessing the power of transformers, researchers have achieved remarkable capabilities in capturing long-range dependencies in sequential data, leading to state-of-the-art performance on many NLP tasks. An exemplary advancement in LLM that has captured the popular consciousness is the development of ChatGPT, a natural language model based on OpenAI’s GPT-3.5/4 architecture.^8^ ChatGPT has garnered attention for its impressive ability to generate coherent and contextually relevant responses across diverse domains, and reached 100 million monthly active users just two months after its launch, making it the fastest-growing (and sustained) consumer application in history.^12^

Additionally, the success of the transformer architecture has extended beyond natural language to other domains, including biology. Protein sequences, which directly encode the complex 3-dimensional structures and functionalities of proteins, have become an intriguing target for LLM research. To address the intricacies associated with protein sequences, researchers have adapted LLM techniques to create protein language models (PLMs) such as ProteinBERT^13^, ProtBERT^14^, ProtT5^14^, and ESM.^15^ These models are specifically designed to capture the intrinsic relationships between amino acids by analyzing sequences at once, enabling tasks such as secondary structure prediction and subcellular location prediction, which act at the token and sequence level respectively.^16^

Numerous studies have explored the potential of fine-tuning PLMs for various downstream tasks, such as engineering protein function^17-19^, identifying sites of post-translational modification^20^, and assessing toxicity.^21^ Beyond prediction of specific outputs, PLMs have begun to build on the remarkable achievements AlphaFold2^22^ in predicting protein 3-dimensional structures. The trRosettaX-Single model^23^ performs similarly to AlphaFold2 without requirement of multiple sequence alignments (slow, CPU-based step to find most similar examples) greatly improving the speed of 3-dimensional structure prediction. However, despite these successes, there has been a notable absence of investigations into the ability of PLMs to effectively learn mutational tasks. We hypothesize this is likely due to the phenomenon that incredibly small changes in sequence (commonly 1 out of 200-400 amino acids) can lead to dramatic changes in structural or functional outcomes.^24^

To evaluate the potential of PLMs in addressing the intricate challenge of learning mutational tasks, we introduce a novel machine learning strategy named “hint token learning” (HTL). HTL is engineered to enhance the ability of PLMs to discern and emphasize the critical impacts of mutations in protein sequences. Our approach is rigorously demonstrated through its superior performance in comparison with conventional fine-tuning methods, as illustrated in three challenging and varied case studies. This innovative strategy effectively tackles some of the primary complexities inherent in mutational datasets, thereby marking a significant advancement in the realm of LLM techniques within computational biology.

## RESULTS

### CASE STUDY 1: PROTEIN STABILITY

Our initial mutational predictive model was centered on Tsuboyama *et al*.’s^25^ comprehensive protein stability dataset, for which we meticulously compiled and pre-processed the data. Table 1 provides a summary of the data, including key metrics such as the total number of data points, the count of unique domains, and the distribution percentages of free energy of unfolding (ΔG) across various bins in all machine learning sets. To evaluate other characteristics of the Tsuboyama dataset, we performed exploratory data analysis (EDA) and investigated the relationship between mutation type and the resultant stabilization or destabilization (Figures, S1-S2 Table S2 and https://github.com/ejp-lab/EJPLab_Computational_Projects/tree/master/HintTokenLearning/EDA). We determined that there were no obvious general trends beyond proline being the most destabilizing type of mutation across the entire dataset (Figure S1). However, when we investigated the effects of mutations of specific amino acids within protein domains, we found diverse and specific trends. For example, Figure S2 shows a bar chart with an amino acid stabilization profile unlike the average. This yeast Myo5 SH3 domain (PDB:1YP5) exhibited strong destabilization by acidic amino acids and an uncharacteristic stabilization by cysteine compared to all other types of mutations. Looking deeper, we tested to see whether certain mutations had greater or lesser effects within different secondary structural motifs. Using DSSP^26^ to generate the secondary structure labels for each wild-type domain (or AlphaFold2 as provided by Tsuboyama *et al*.), we computed the average effect of mutations in secondary structures both generally and in each specific domain. Here, we identified the same phenomenon that was observed with the mutation types. Generally, there does not appear to be a particular secondary structural motif sensitive to mutation, beyond B (residue in isolated β-bridge) or I (5-turn helix) type secondary structures exhibiting small destabilizing preferences (Figure S3). However, within domains (based on present motifs), there are unique and specific trends. For example, Figure S4 displays an example of a multi secondary structured domain (1EKL) which exhibits preferential stabilization and destabilization. This type III antifreeze protein domain is more strongly stabilized by mutations in G (3-turn helix) and T (hydrogen bonded turn) type secondary structure while simultaneously being significantly destabilized by mutations in B, E (extended strand in parallel and/or anti-parallel β-sheet), H (4-turn helix), and S (bend) secondary structure. Together, these analyses support the idea that simple and mutation site localized heuristics are insufficient for predicting effects on protein stability.

**Table 1.**
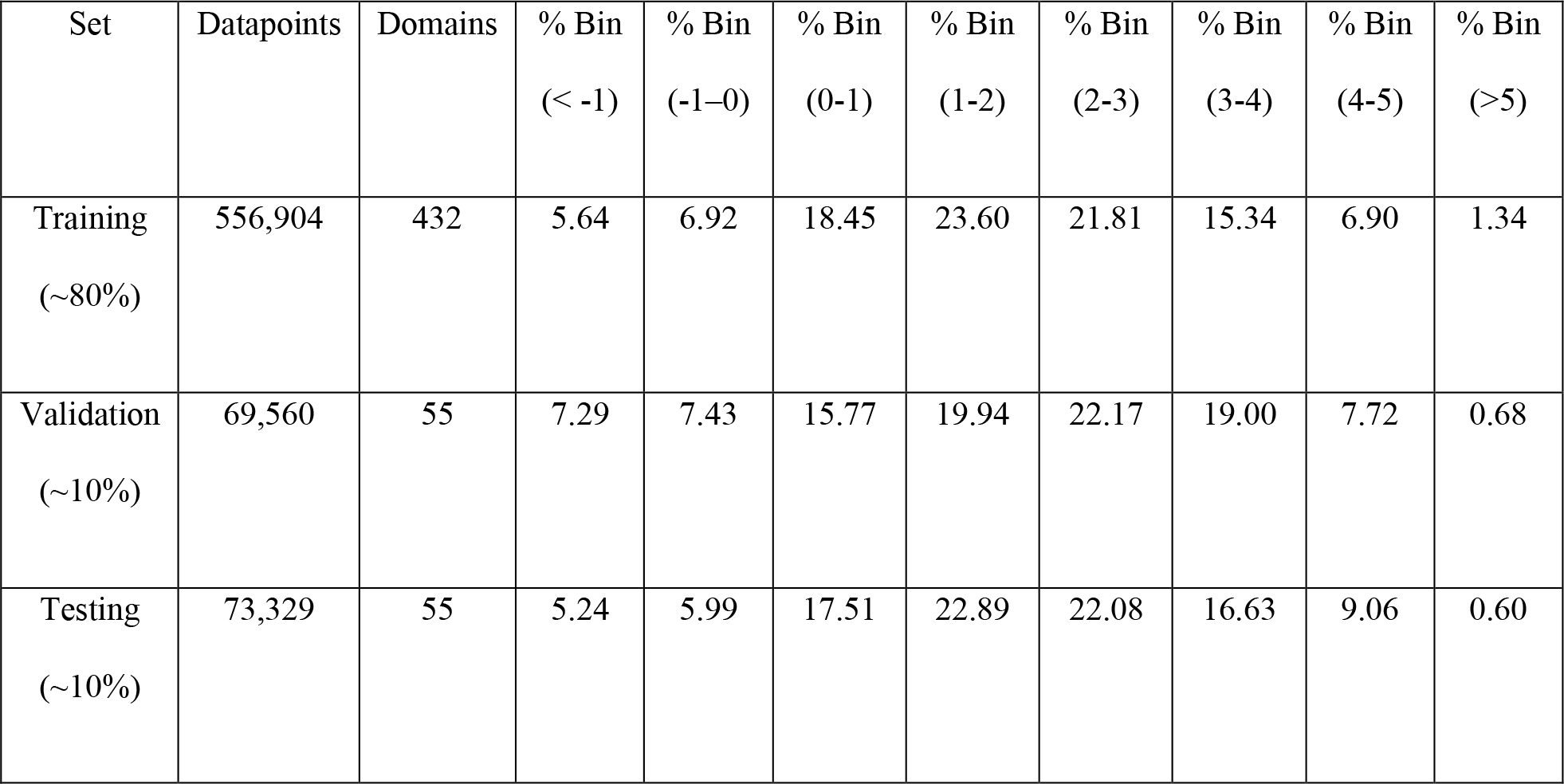
Statistics of machine learning sets derived from Tsuboyama protein stability dataset.

| Set | Datapoints | Domains | % Bin<br>(< -1) | % Bin<br>(-1-0) | % Bin<br>(0-1) | % Bin<br>(1-2) | % Bin<br>(2-3) | % Bin<br>(3-4) | % Bin<br>(4-5) | % Bin<br>(>5) |
| --- | --- | --- | --- | --- | --- | --- | --- | --- | --- | --- |
| Training<br>(~80%) | 556,904 | 432 | 5.64 | 6.92 | 18.45 | 23.60 | 21.81 | 15.34 | 6.90 | 1.34 |
| Validation<br>(~10%) | 69,560 | 55 | 7.29 | 7.43 | 15.77 | 19.94 | 22.17 | 19.00 | 7.72 | 0.68 |
| Testing<br>(~10%) | 73,329 | 55 | 5.24 | 5.99 | 17.51 | 22.89 | 22.08 | 16.63 | 9.06 | 0.60 |

We therefore wanted to test the hypothesis that finetuning a PLM, which can analyze entire sequences at once, could effectively learn how mutations alter protein folding stability. To do this, we finetuned the ProtBERT model (a lighter-weight PLM accessible to most researchers) on the Tsuboyama dataset, starting with a conservative set of hyperparameters as a scouting run. To our surprise, we observed a very poor ability to regress the Tsuboyama dataset (R: ∼0.1). We recognized that even with improvements from hypertuning we would need to devise a new solution to improve the quality of this model. Our resulting strategy was to train a model with the same hyperparameters, but this time on sequences which were tokenized with the two new special tokens (MUT_start and MUT_end) surrounding the mutation sites to help the model focus on these changes (“hint token learning”, Figure 1).

**Figure 1.**
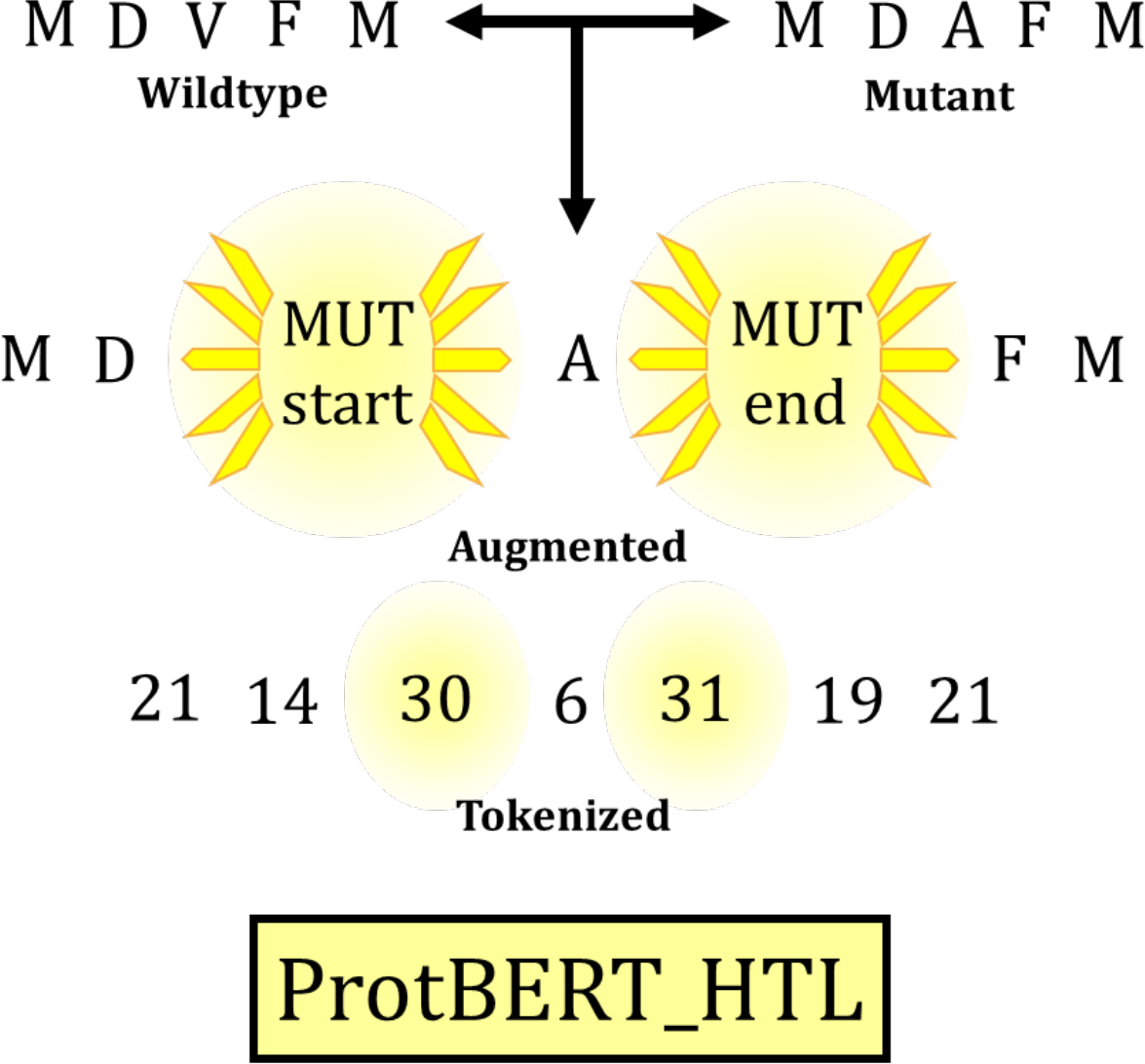
Diagram of the hint token learning process.

Immediately, we observed a significant improvement in the ability to regress the Tsuboyama dataset (R^2^: ∼0.4). Both the standard ProtBERT model (ProtBERT_ΔG) and the hint token learning ProtBERT model (ProtHTL_ΔG) were tuned to produce optimal models which could be evaluated on the testing set. Bar graphs displaying the testing set mean squared error (MSE) values per bin and in total for these two models are displayed in Figure 2.

**Figure 2.**
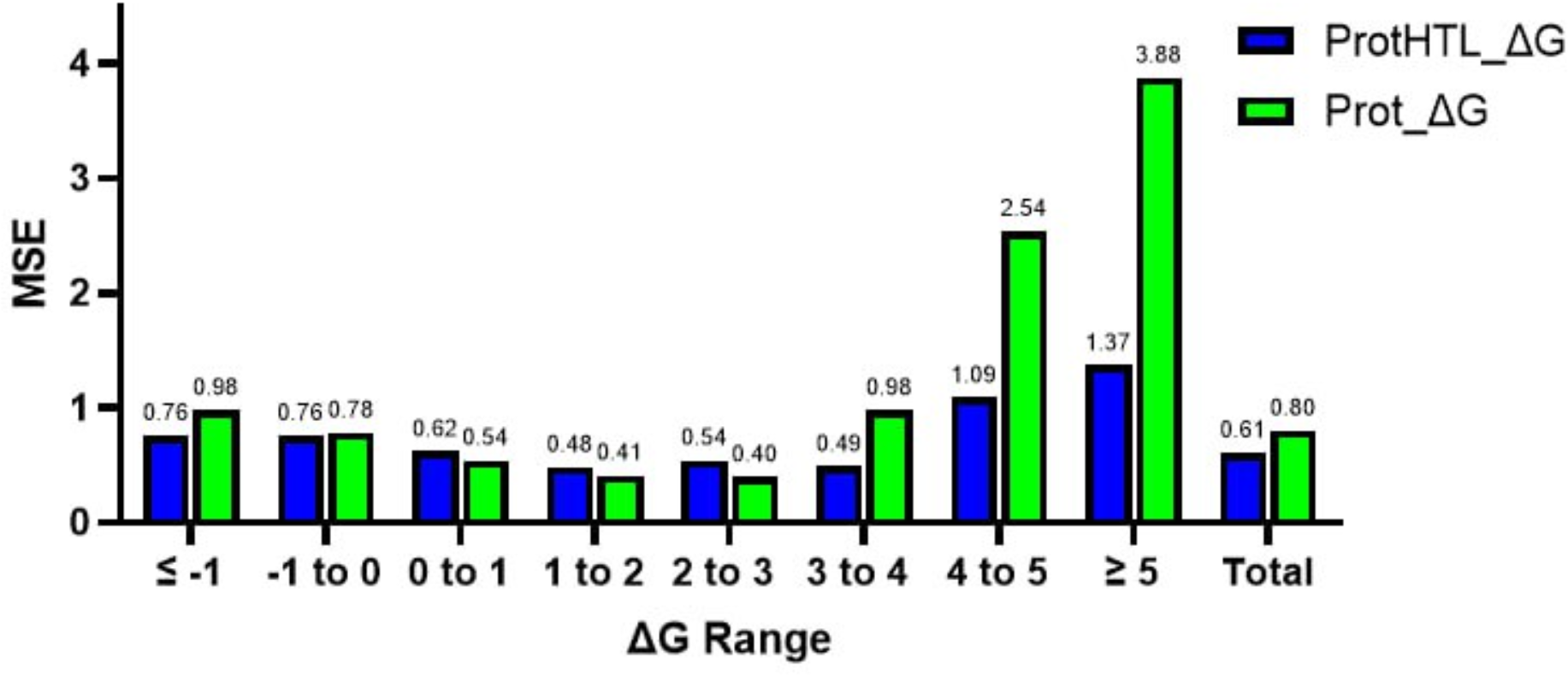
Bar graph contrasting the Prot_ΔG (green) and ProtHTL_ΔG (blue) models in each bin and overall, on the testing set. Bin range units are in kcal/mol.

Binned error analysis demonstrated that the ProtHTL_ΔG made predictions more accurately than Prot_ΔG in almost all cases across the range of the target variable, but most significantly in the strongly stabilizing mutations (values 3 kcal/mol and above). Overall, the ProtHTL_ΔG produced a MSE of ∼23.75% lower than Prot_ΔG in general, but ∼54.8% lower than Prot_ΔG MSE for highly stabilizing mutations. ProtHTL_ΔG also performed better on the most destabilizing mutations, exhibiting ∼22.5% MSE lower than Prot_ΔG. This is easily explained by the propensity of Prot_ΔG to predict with dummy like regression-like behavior shown in Figure 3.

**Figure 3.**
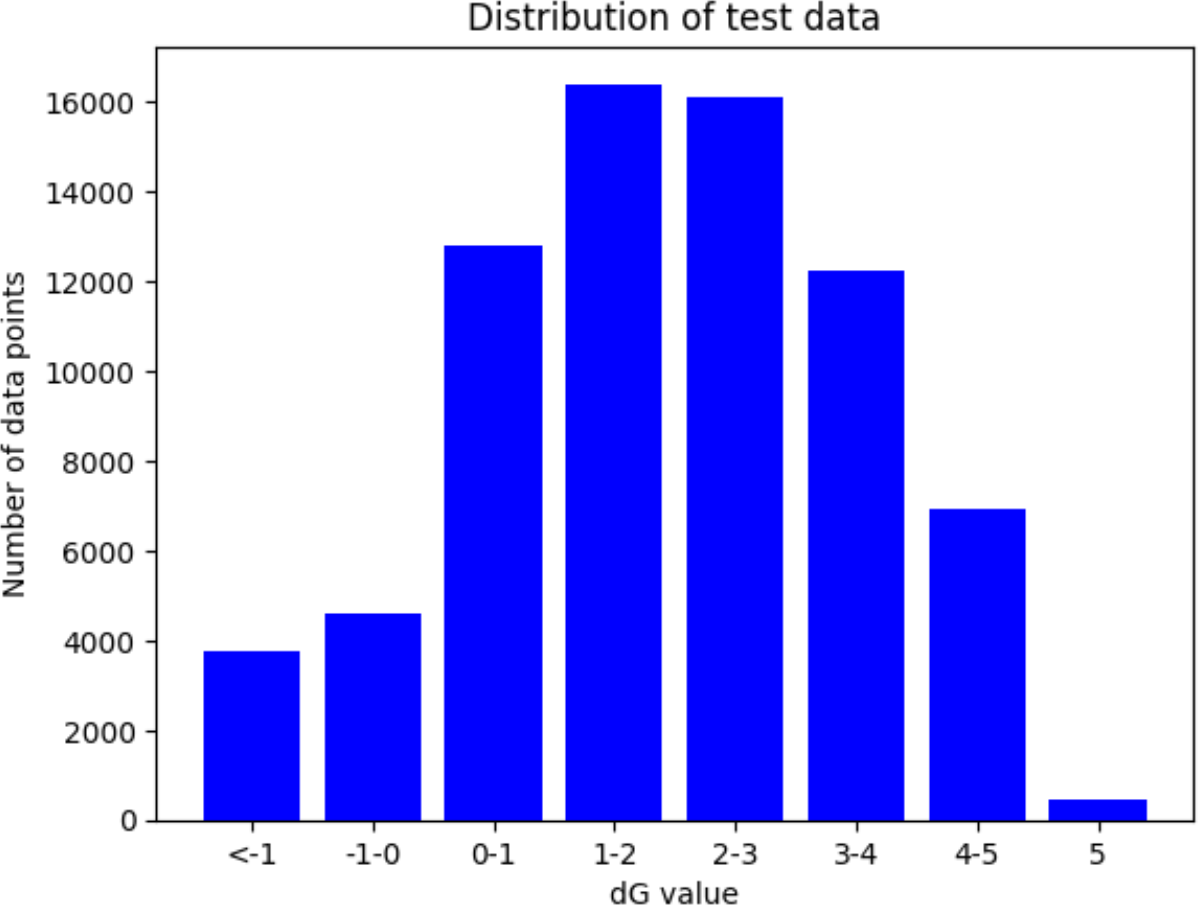
Bar plot showing the number of predictions in each bin.

### CASE STUDY 2: GFP BRIGHTNESS

In our previous case study, we noted substantial improvements in protein stability models (Prot_ΔG and ProtHTL_ΔG) using the HTL approach. To determine if this is a broader phenomenon, we applied both traditional and HTL fine-tuning methods to the Sarkisyan^27^ green fluorescent protein (GFP) mutant brightness dataset. Additionally, due to the large amount of available data on the single GFP protein, we experimented the effect of titrating the amount of training data available for fine-tuning. Prot_GFP (pretrained ProtBERT model) and ProtHTL_GFP (pretrained ProtBERT model utilizing hint tokenized GFP mutant sequences) were fine-tuned on the Sarkisyan dataset and the outcomes are detailed in Table 2. These data reveal that regardless of the training data volume (1%, 10%, or 80%), the HTL model consistently yielded more accurate predictions of mutant GFP brightness than the traditional fine-tuning method (first two data columns). Specifically, at 1% training data volume, we observed a near 100,000-fold increase in R^2^, although both predictions are admittedly poor in an absolute sense. While the R^2^ increases at 10% and 80% training volumes were less dramatic compared to the 1% volume, they were still notable, with increases of 8.57% and 10.13%, respectively.

**Table 2.** Benchmarking HTL on independent testing set vs traditional finetuning at various amounts of GFP training data.

|  | Prot_GFP | ProtHTL_GFP | Prot_ΔG_GFP | ProtHTL_ΔG_GFP |
| --- | --- | --- | --- | --- |
| Pretrained model | ProtBERT | ProtBERT | Prot_ΔG | ProtHTL_ΔG |
| Finetuning method | Traditional | HTL | Traditional | HTL |
| Training Data (%) | ($R^2$) | ($R^2$) | ($R^2$) | ($R^2$) |
| 1 | 3.58 E-05 | 4.90 E-02 | 0.23 | 0.23 |
| 10 | 0.35 | 0.38 | 0.58 | 0.58 |
| 80 | 0.79 | 0.87 | 0.84 | 0.84 |

While HTL realized greater performance over traditional fine-tuning at lower amounts of training data (representative of real-life laboratory situations), we sought even better performing models. Knowing that the brightness of GFP is in part related to the stability of the chromophore, we attempted to fine-tune the Sarkisyan dataset starting from the now pretrained Prot_ΔG and ProtHTL_ΔG models from case study 1 (Table 2, last 2 data columns).

Encouragingly, beginning with either of the protein stability models, Prot_ΔG or ProtHTL_ΔG, significantly enhanced our ability to predict mutant GFP brightness, particularly at lower training data volumes. At 1% training volume, utilizing either of the protein stability models yielded a R^2^ % increase of ∼370%. This case study underscores the efficacy and practical value of transfer learning, especially when leveraging a robust HTL-based protein stability model. While the use of HTL upon the protein stability models did not appear to provide an additional benefit in predictive accuracy in the transfer learning scenario, we note that the highest accuracy (R^2^ = 0.87) was achieved with Prot_HTL_GFP with 80% training data.

### CASE STUDY 3: RECA BIOACTIVTY

The final case study we utilized to evaluate the power of HTL was a RecA bioactivity dataset compiled from McGrew *et al*.^28^ This dataset includes nearly 1000 mutants of the RecA protein with an accompanying activity label. These labels were used to make a three-part classifier with activities greater than or equal to wild-type (rec+), similar, but less than wild-type (rec±), and no activity (rec-). As in case study 2, we trained 4 models: Prot_RecA, ProtHTL_RecA, Prot_ΔG_RecA, and ProtHTL_ΔG_RecA to test the effectiveness of HTL and to further benchmark the transfer learning capabilities of the protein stability models. Figure 4 shows plotted confusion matrices for all four models.

**Figure 4.**
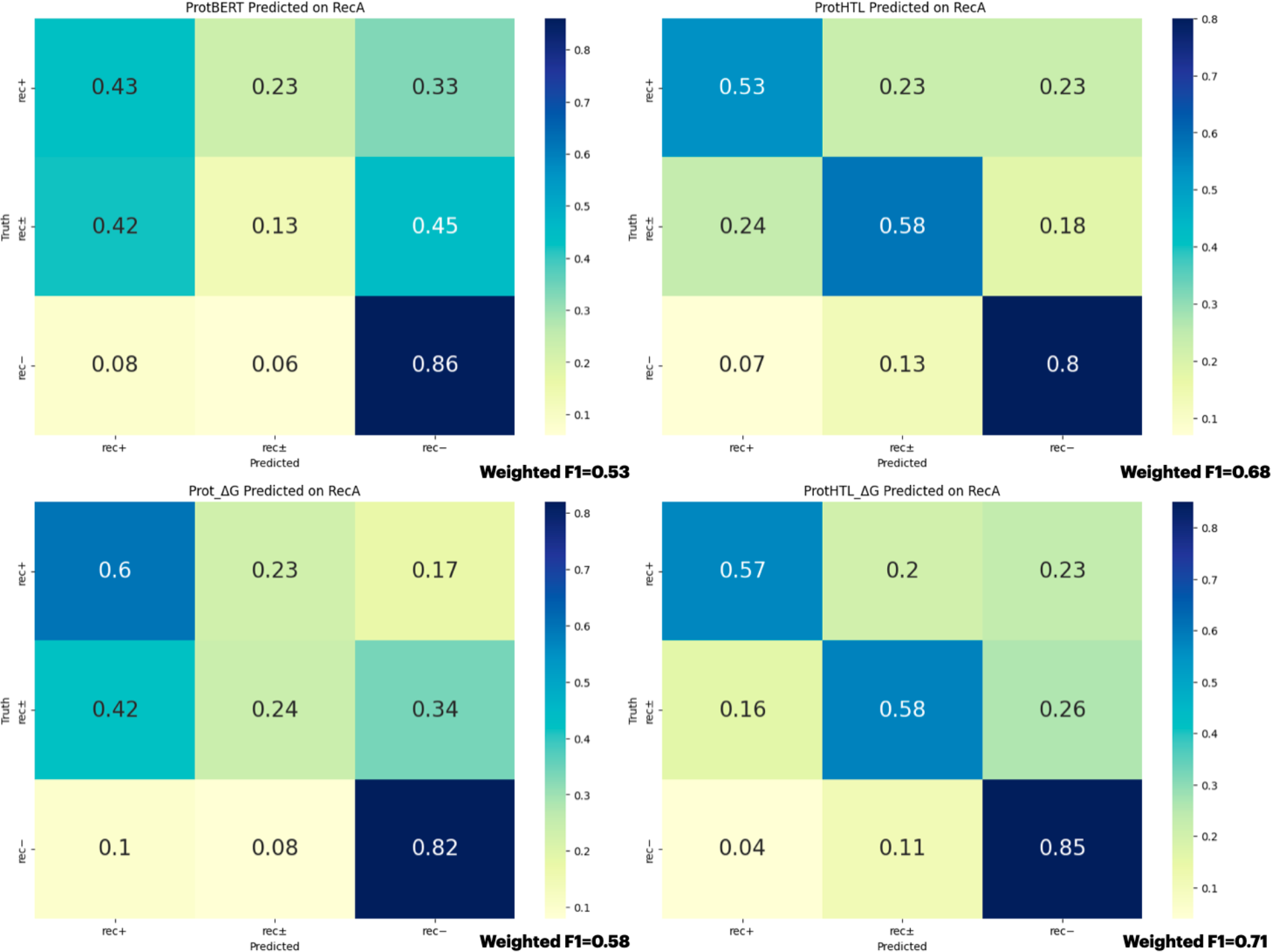
Confusion matrices of independent testing set of RecA case study. The top row displays the performance of Prot_RecA (left) vs ProtHTL_RecA (right). The bottom row displays the performance of Prot_ΔG_RecA (left) and ProtHTL_ΔG_RecA (right). The left column demonstrates the change in performance using the same finetuning strategy but changing the pretrained model from ProtBERT (top) to case study 1 protein stability model (bottom). Cooler colors on the color bar indicate larger values.

The RecA investigation demonstrated that HTL was effective in improving the F1 score of the independent testing set over the traditional fine-tuning method in the case of the ProtBERT pretrained models. Use of HTL was able to improve the weighted F1 score of the independent test set by an astounding 28.30%. Furthermore, as was seen in the case of the GFP case study, the RecA bioactivity prediction was aided by using one of the protein stability models as the pretrained model over ProtBERT itself. Independent of HTL, use of the protein stability model as the pretrained model yielded a weighted F1 increase of 9.43%. Furthermore, we also observed that HTL improved the predictive power of the fine-tuning of the protein stability model ProtHTL_ΔG by an additional 4.41%. These findings unambiguously show the efficacy of HTL and demonstrate the versatility of transfer learning from protein stability models in various downstream applications.

### EXPLANATION OF THE HINT TOKEN

We investigated our models to potentially glean ways in which HTL aids in improving model predictivity. To do this, we produced attention plots for the protein stability independent testing dataset. By analyzing the attention weights at each node in the sequences between Prot_ΔG and ProtHTL_ΔG, we can see where focus is shifting as a function of the incorporation of the hint token. Somewhat counterintuitively, we observed that hint tokens do not act exclusively as positional flags, but rather that attention is very often shifted to other residues in the sequence compared to the no hint token control. Additionally, it is worth noting that the hint tokens themselves do not receive a disproportionate amount of attention from the model. Across the dataset we observed that total attention to hint tokens for single mutant datapoints was on average ∼5%. A representative example of the attention shifting phenomenon is shown in Figure 5.

**Figure 5.**
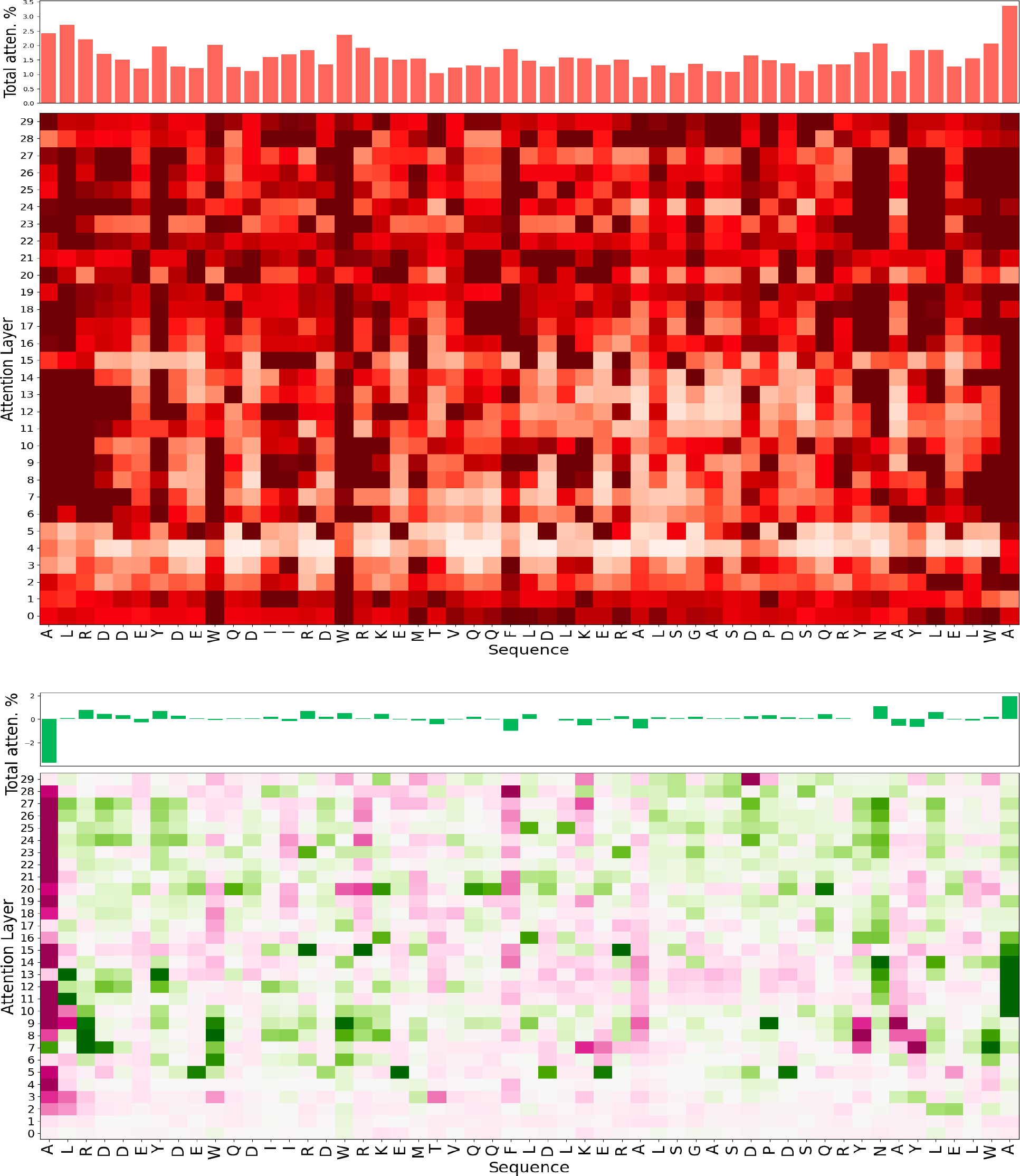
Attention plots for a representative testing dataset example sequence. The top figure displays the visualized attention weights from the ProtHTL_ΔG model omitting weight contribution from the hint tokens themselves. Darker coloration indicates greater attention. The bottom figure displays the change in attention weights between all real sequence positions from ProtHTL_ΔG and Prot_ΔG omitting weight contribution from the hint tokens themselves. Green coloring indicates an increase in attention at the sequence position from incorporation of the hint tokens. Conversely, purple indicates a loss of attention as a function of incorporation of hint tokens.

In this example sequence, we see that for the ProtHTL_ΔG model (top portion) that attention is somewhat uniform across the sequence and that total attention at the hint tokens is approximately 4%. However, when comparing the change in attention at the real sequence positions (ProtHTL_ΔG - Prot_ΔG) we observe a significant decrease in attention at the far N-terminal region of the sequence and two significant increases in attention directly surrounding the mutation site and approximately 12-16 residues N-terminal to the mutation site. Additionally, we visualized the embeddings of our protein stability models.

## DISCUSSION

This study aimed to enhance the ability of protein language modeling towards accurate prediction of mutational datasets. By introducing the HTL strategy, we were able to successfully train a highly accurate ΔG of folding model (ProtHTL_ΔG) with superior accuracy to a traditional pretrained PLM (Prot_ΔG). The promise of ProtHTL_ΔG is especially exciting as it effectively predicted the tails of the stability distribution while Prot_ΔG tended to over-predict towards the mean.

Investigation of HTL through attention plots and embedding analysis strongly supported the hypothesis that PLMs are suitable for studying mutational datasets. Despite the similarity between entries in mutational datasets, PLMs have the advantage of observing the entire sequence at once, unlike local calculations or heuristics. We observed that the introduction of hint tokens into sequences shifted the model’s focus to different areas of the sequence and resulted in distinct model embeddings. ProtHTL_ΔG serves as an example of how PLMs aided by HTL can capture the complex relationships between amino acids in sequences and effectively combat the challenges presented by small changes in protein sequence leading to large phenotype perturbations.

Principally, we believe that access to an accurate protein stability model, especially one with the capability to make accurate predictions at the extremes of stability, will be instrumental in studying many of the outstanding problems in biology and biologics discovery. In particular, we see ProtHTL_ΔG leading to major advancements in the understanding and design of protein folds, with broad applications to protein engineering, where changes in structure to escape local minima are inhibited due to poor predictability on the edges of the known sequence landscape. For instance, ProtHTL_ΔG could be used within the Rosetta design framework for optimizing antibody or enzyme stability.

Beyond hint token learning, the versatility of the ProtHTL_ΔG model was performance tested by its application to several downstream tasks. Most excitingly, ProtHTL_ΔG was not just effective in predicting the brightness of mutations in GFP, which is strongly related to protein stability, but also effective in predicting RecA mutant bioactivity. These transfer learning experiments raised several interesting ideas regarding the limitations of pretrained PLMs. By their very nature, traditional pretrained PLMs are derived from masked language modeling on huge corpora of protein sequences.^29^ This makes them distinctly effective at capturing and recapitulating evolutionary information. However, as discussed Tsuboyama *et al*., evolution and stability are partially at odds. Structures which are overly stable are not in the suitable energy landscape for biological activity. This leads one to question whether pretrained PLMs like ProtBERT are the right starting point for design tasks. We believe that the GFP training data titration experiments where the protein stability models substantially outperformed the pretrained PLMs with low amounts of data are a strong indicator that PLMs are not optimal for design tasks.

We have made ProtHTL_ΔG available within the HuggingFace framework to enhance its accessibility and facilitate its integration into various downstream fine-tuning applications related to energetic phenomena and in protein design. Ultimately, we hope that the introduction of the HTL strategy paves the way for innovative strategies that are waiting to be uncovered.

## METHODS

### DATA PREPROCESSING: PROTEIN STABILITY

The dataset used for training ProtHTL_ΔG is derived from Tsuboyama *et al*.^25^ This dataset is a mega-scale evaluation of protein mutant ΔG of folding and represents the largest database of protein mutant stability to date. The Tsuboyama dataset includes 851,552 high quality data points made from over 1.8 million measurements of protein folding stability. These measurements come from single and double mutations in 542 different protein domains, each of which is between 40-72 residues in length. However, the Tsuboyama dataset also included several mutations which were insertions or deletions. These types of mutations represent a significantly different meaning from the majority of the data and were therefore removed. Additionally, given the nature of their assay with respect to construct transcription and translation from oligonucleotide pools, some of the single or double mutations were not incorporated. For example, in an experiment attempting to measure the double mutant D26E_D28R, where the resultant construct made was actually D26E_D28D, only the first of the two mutations were successfully incorporated. Datapoints like these were also removed, leaving 699,793 datapoints for ML.

The culled Tsuboyama dataset was then split into ∼80-10-10 training, validation, and testing sets where members of each set were kept in groups according to protein domain. Splitting by groups ensures that model training and evaluation is being assessed on out of group generalizability. The explicit enumeration of our datasets can be found on our GitHub at https://github.com/ejp-lab/EJPLab_Computational_Projects/tree/master/HintTokenLearning.

### DATA PREPROCESSING: GFP BRIGTHNESS

The data for training ProtHTL_GFP was derived from Sarkisyan *et al*.^27^ In a procedure similar to preprocessing of the Tsuboyama dataset, the Sarkisyan dataset was culled to only keep single and double mutants of GFP. Curation of splits for ML were made in the following way. For values of 1%, 10%, and 80% of training data, each were further segmented to use 20% as validation data. The remaining data was set aside for testing set evaluation. When datasets were split, they were done with a residue position grouping method. Splitting by position allows there to be no redundant mutation positions between sets and again forces model training and evaluation to assess out-of-group generalizability.

### DATA PREPROCESSING: RECA BIOACTIVTY

The McGrew *et al*. dataset^28^ was processed for ML by creating training, validation, and testing sets through multi-label stratification. Given that this set did not contain saturation data and was for a single protein, we ensured that the number of mutations and the class label distribution was preserved across sets. Due to the smaller overall dataset size, we wanted to ensure that a rigorous and challenging independent dataset for benchmarking. Therefore, we used multi-label stratification to produce 66-17-17 sets for learning.

### HINT TOKENIZATION

Sequences from all datasets were edited to include a MUT_start and MUT_end token around all of the mutations relative to the wild-type construct. For example, the mutation D26E would appear as [MUT_start] E [MUT_end] at position 26 in the sequence. With these modifications made to the culled sequences, the BertTokenizer was imported from the transformers Python library and instantiated with from_pretrained method and the following arguments: Rostlab/prot_bert, do_lower_case=False. This base ProtBERT tokenizer was then updated to include the MUT_start and MUT_end tokens with the add_tokens method. Note that for the wild-type control sequences, no MUT_start or MUT_end tokens were incorporated.

### MODEL TRAINING

We trained the ProtHTL_ΔG regression model using the Trainer class from the transformers Python library. Training arguments for the Trainer object included torch_compile=True, bf16=True, and optim=adamw_torch_fused to make use of PyTorch 2.0.^30^ The final models from each case study were selected from the best trial from Bayesian hyperparameter searching with the TPE sampler of the Optuna Python library. Fifty trials were searched while attempting to minimize the mean squared error loss of the validation set for regression problems, and cross entropy loss for classification tasks. Models were trained for a maximum of 10 epochs with an early stopping patience of 5 epochs. Finally, the tuned models were used to predict the data of the held-out testing set.

### HINT TOKEN ATTENTION PLOTING

In order to better understand the effect of the hint tokens on our models, we visualized model attention weights in attention plots. We visualized these plots in two distinct ways utilizing weights from both the ProtHTL_ΔG and Prot_ΔG models. These models employ 30 attention layers each with 16 heads. For each layer, the weights for all 16 attention heads were summed. The bar graph portion of the hint token learning attention plot (top in Figure 5) displays the % attention for each token in the sequence. The heatmap beneath displays the attention weight in each of the 30 attention layers at each position in the sequence. These plots are highlighted such that darker shades of red show increased attention. The delta attention plot (bottom in Figure 5) is a similar analysis but shows the attentions at positions in the ProtHTL_ΔG model subtracted by the Prot_ΔG model. This plot omits the positions of the hint tokens for clarity. Finally, these plots are colored such that green coloring indicates an increase in attention while purple indicates a loss of attention.

## DATA AVAILABILITY

The data used to train ProtHTL_ΔG and ProtHTL_GFP including all training, validation, and testing splits are available on our GitHub at https://github.com/ejp-lab/EJPLab_Computational_Projects/tree/master/HintTokenLearning/Data. Additionally, the raw data from Tsuboyama *et al*. is available at https://doi.org/10.5281/zenodo.7844779.

## CODE AVAILABILITY

The code for hint token learning and a guide to reproducing the results demonstrated herein is available on our GitHub at https://github.com/ejp-lab/EJPLab_Computational_Projects/tree/master/HintTokenLearning.

## ACKNOWLEDGMENTS

This work was supported by the University of Pennsylvania and the National Institutes of Health (NIH R01-GM127593 to E.J.P.). S.G.G. thanks the National Science Foundation (NSF) for funding through the NSF Graduate Research Fellowship Program (DGE-1845298). R.M.P. thanks the NIH for funding through the Chemistry Biology Interface Training Program (T32-GM133398).

## AUTHOR INFORMATION

* SGG; EJP: X.L, R.P, and S.G.G each performed data preprocessing, model training, and data analysis guided by EJP. The manuscript was written through contributions of all authors. All authors have given approval to the final version of the manuscript.

## ETHICS DECLARATIONS

The authors declare no competing interests.

