## Supporting Information for "Proteins Need Extra Attention: Improving the Predictive Power of Protein Language Models on Mutational Datasets with Hint Tokens"

**Electronic Supplementary Information**

**Table of Contents:**

1. **Software…………………………………………………………………………………. S2**
2. **Datasets…………………………………………………………………………………. S2**
3. **EDA……….……………………………………………………………………............... S2**
4. **Hint Token Investigation……………………………………………………………… S15**
5. **Model Training………………………..……………………………………………….. S22**
6. **Reference…..……………………………………………………………………………… S25**

### Software

The software used for this work can be found in the conda environment yml file alongside installation instructions at the following link:

https://github.com/ejp-lab/EJPLab_Computational_Projects/tree/master/HintTokenLearning/Anaconda.

### Datasets

The Tsuboyama dataset was acquired from the following link. https://zenodo.org/record/7401275.

The Sarkisyan dataset was acquired from the following link. https://static-content.springer.com/esm/art%3A10.1038%2Fs41467-023-38099-z/MediaObjects/41467_2023_38099_MOESM13_ESM.xlsx The RecA dataset was extracted from the McGrew review paper[1].

All machine learning datasets were created as described in the main text. For reproducibility, these datasets can be found at our GitHub through the following link: https://github.com/ejp-lab/EJPLab_Computational_Projects/tree/master/HintTokenLearning/Data.

### EDA

The Tsuboyama dataset was explored thoroughly for potential heuristics which explain stabilization of protein domains. Firstly, we wrote a Python script which found the average effect of stabilization/destabilization from mutation as a function of amino acid type. Figure S1 displays a bar chart for each unique amino acid mutation type in the dataset.

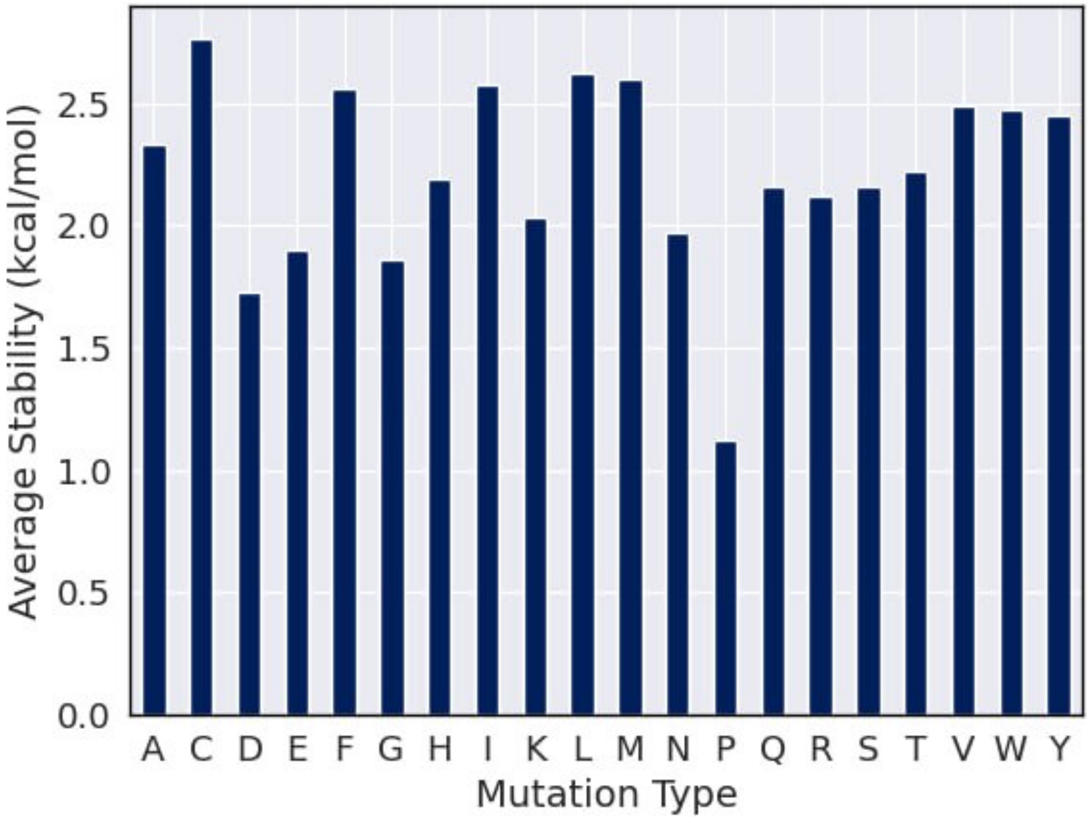


**Figure S1.** Bar chart plotting average stability in kcal/mol per mutation type in the Tsuboyama dataset.

Seeing as this global analysis largely only provided insight that proline appears as the most destabilizing of the types of mutations, we investigated each of the protein domains individually. Here we found much more informative trends. Figure S2 presents an example bar chart (yeast Myo5 SH3 domain) where a unique stabilization/destabilization profile is observed relative to the average of the set. The csv titled DomainStatisticsPerMutationType.csv on our GitHub at https://github.com/ejp-lab/EJPLab_Computational_Projects/tree/master/HintTokenLearning/EDA displays the average stabilization/destabilization values for all mutation types in every domain. For clarity, Table S1 provides relevant statistics of this csv file.

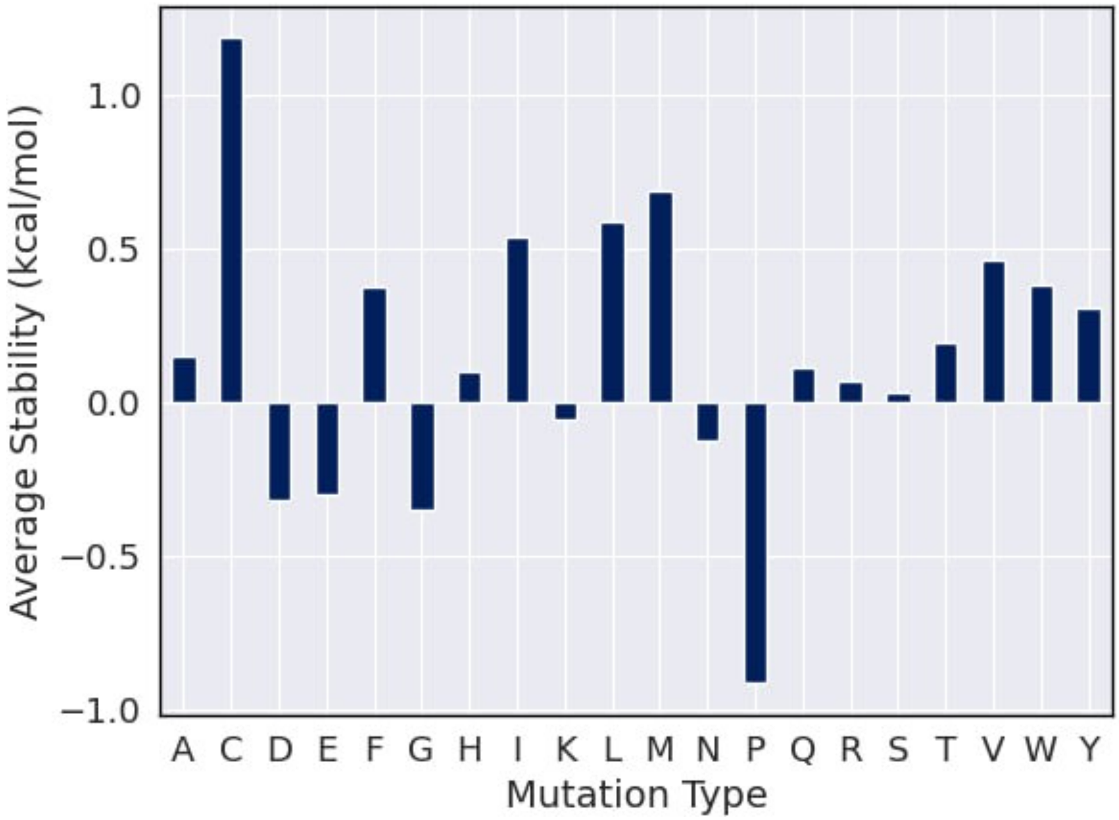


**Figure S2.** Bar chart plotting average stability in kcal/mol per mutation type in a yeast Myo5 SH3 domain (1YP5).

**Table S1.** Table displaying statistics of mutation types on the Tsuboyama dataset.

| Mutation Type | Minimum | Maximum | Mean | Stdev |
|---|---|---|---|---|
| A | -0.968 | 4.71 | 2.314 | 1.025 |
| C | 0.768 | 4.97 | 2.751 | 0.843 |
| D | -0.874 | 4.187 | 1.698 | 0.92 |

| | | | | |
|---|---|---|---|---|
| E | -0.81 | 4.252 | 1.875 | 0.929 |
| F | 0.342 | 4.837 | 2.554 | 0.963 |
| G | -0.496 | 4.747 | 1.842 | 1.001 |
| H | -0.294 | 4.558 | 2.158 | 0.985 |
| I | -0.256 | 4.859 | 2.586 | 0.954 |
| K | -0.906 | 4.491 | 1.977 | 0.94 |
| L | -0.938 | 4.851 | 2.634 | 0.985 |
| M | 0.288 | 4.976 | 2.616 | 0.984 |
| N | -0.469 | 4.655 | 1.946 | 0.98 |
| P | -0.912 | 3.898 | 1.094 | 0.917 |
| Q | -0.395 | 4.515 | 2.12 | 0.957 |
| R | -0.045 | 4.463 | 2.094 | 0.887 |
| S | -0.396 | 4.837 | 2.139 | 0.997 |
| T | -0.043 | 4.716 | 2.213 | 1.006 |
| V | 0.203 | 4.917 | 2.503 | 0.981 |
| W | 0.372 | 4.785 | 2.465 | 0.923 |
| Y | 0.308 | 4.78 | 2.439 | 0.95 |

The same analyses were performed for protein domain secondary structure motifs. Figure S3 displays a bar chart of the global dataset metrics, while Figure S4 shows a unique, representative of a specific domain. Again, we observed that the global analysis was mostly uninformative, but that individual protein domain analysis reveals many specific trends. Stabilization data for each WT fold can be found at The csv titled DomainStatisticsPerMutationType.csv on our GitHub at https://github.com/ejp-lab/EJPLab_Computational_Projects/tree/master/HintTokenLearning/EDA, but representative metrics for this datasheet can be found in Table S2.

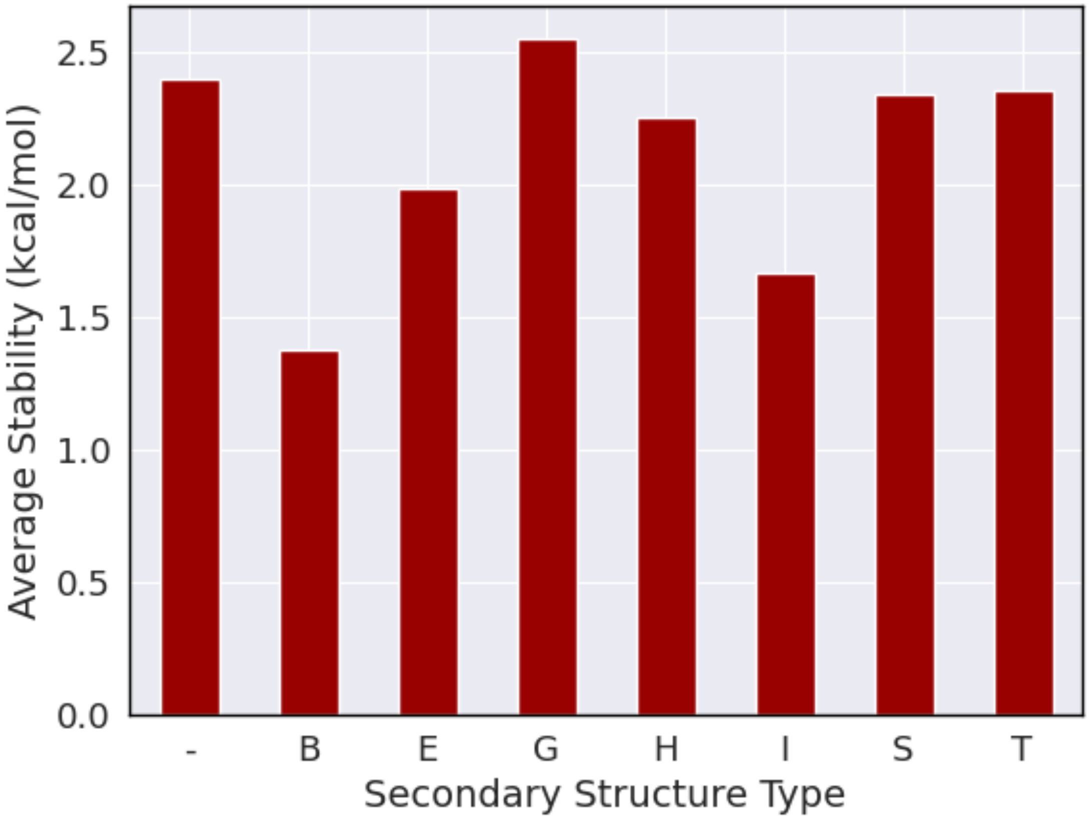


**Figure S3.** Bar chart plotting average stability in kcal/mol per secondary structure type from.

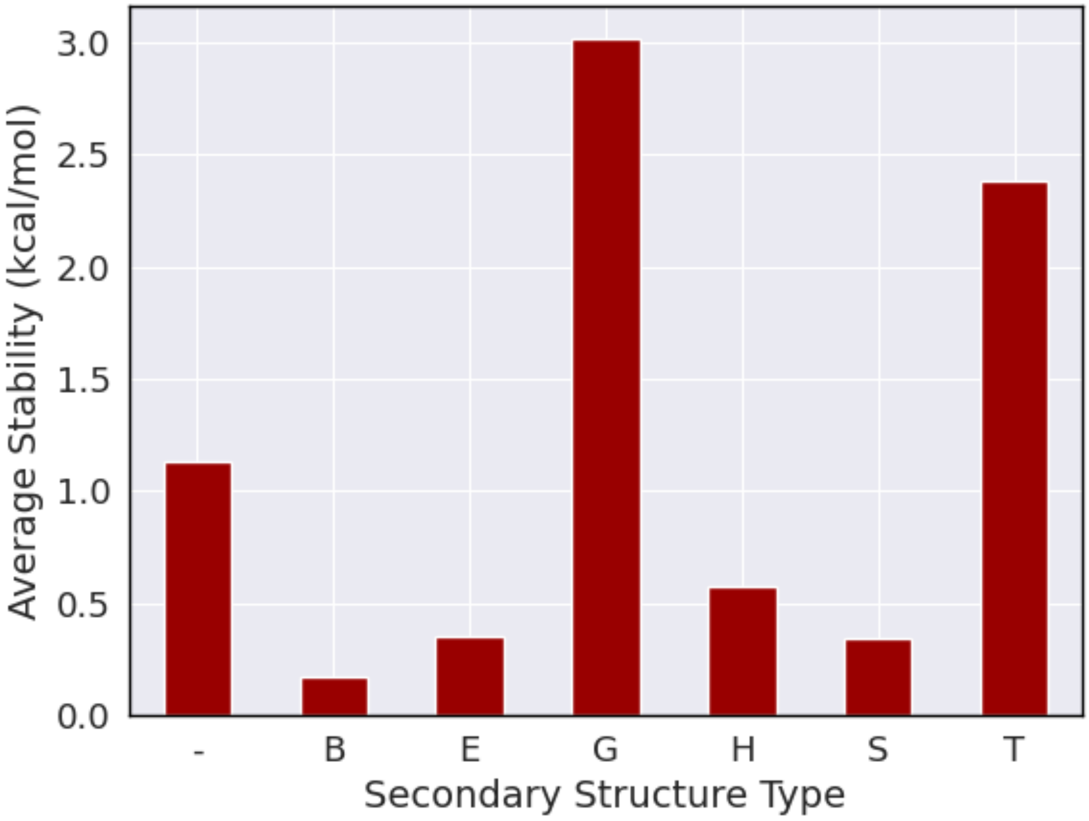


**Figure S4.** Bar chart plotting stability in kcal/mol per secondary in a type III antifreeze protein (1EKL).

**Table S2.** Table displaying statistics of secondary structure on the Tsuboyama dataset.

| Secondary Structure | Minimum | Maximum | Mean | Stdev |
|---|---|---|---|---|
| - | 0.277 | 4.765 | 2.365 | 0.96 |
| B | -0.315 | 4.419 | 1.446 | 1.19 |
| E | -0.106 | 4.61 | 1.711 | 1.205 |
| G | -0.255 | 4.764 | 2.264 | 1.001 |

| | | | | |
|---|---|---|---|---|
| H | -0.477 | 4.947 | 1.882 | 1.269 |
| I | -0.289 | 4.847 | 0.309 | 0.767 |
| S | -0.698 | 4.686 | 0.79 | 1.255 |
| T | 0 | 1.668 | 0.004 | 0.078 |

We conducted similar EDA for GFP protein to explain the relationship between brightness and residual mutations. Similarly with the analysis of Tsuboyama dataset, we wrote a Python script which calculated the average effect of protein brightness change from mutation as a function of amino acid type. Figure S5 displays a bar chart for each unique amino acid mutation type in the dataset and its corresponding brightness.

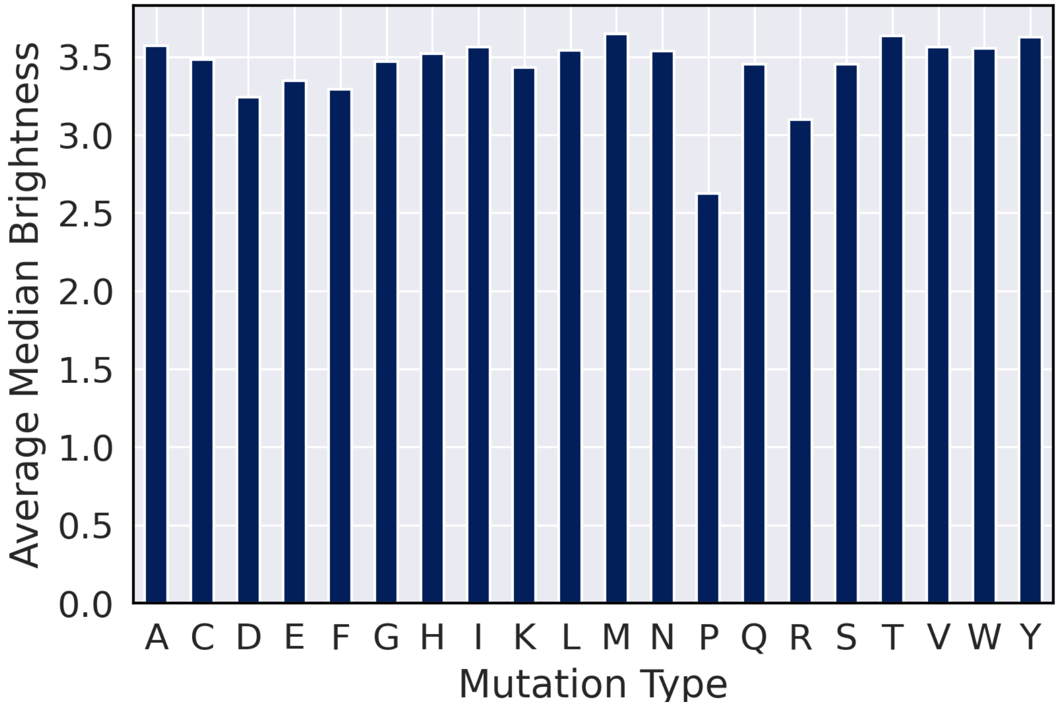


**Figure S5**. Bar chart plotting average median brightness per mutation type in the Sarkisyan dataset.

This bar plot shows that proline appears as the most influential type of mutation on GFP brightness. To give a better insight into the general trend of brightness as the effect of each mutation type, we provide relevant statistics in Table S3.

**Table S3.** Table displaying statistics of mutation types on Sarkisyan dataset.

| Mutation Type | Minimum | Maximum | Mean | Stdev |
|---|---|---|---|---|
| A | 1.301 | 3.833 | 3.572 | 0.295 |
| C | 1.301 | 3.855 | 3.482 | 0.572 |
| D | 1.301 | 3.771 | 3.241 | 0.845 |
| E | 1.301 | 3.874 | 3.345 | 0.769 |
| F | 1.299 | 3.786 | 3.293 | 0.798 |
| G | 1.301 | 4.114 | 3.471 | 0.511 |
| H | 1.301 | 3.895 | 3.520 | 0.461 |
| I | 1.395 | 3.834 | 3.564 | 0.332 |
| K | 1.301 | 3.746 | 3.431 | 0.678 |
| L | 1.301 | 3.992 | 3.541 | 0.417 |
| M | 3.135 | 3.829 | 3.647 | 0.140 |
| N | 1.301 | 3.863 | 3.539 | 0.446 |
| P | 1.301 | 3.785 | 2.626 | 1.034 |

| | | | | |
|---|---|---|---|---|
| Q | 1.301 | 3.837 | 3.452 | 0.625 |
| R | 1.301 | 3.785 | 3.097 | 0.976 |
| S | 1.301 | 3.873 | 3.452 | 0.584 |
| T | 3.122 | 4.008 | 3.635 | 0.166 |
| V | 1.301 | 3.956 | 3.563 | 0.451 |
| W | 3.380 | 3.694 | 3.553 | 0.159 |
| Y | 3.127 | 3.838 | 3.625 | 0.142 |

The same analyses were performed for secondary structure motifs. Figure S6 displays a bar chart of the average brightness per secondary structure type. In addition, we provide statistics of brightness for secondary structure motifs in Table S4.

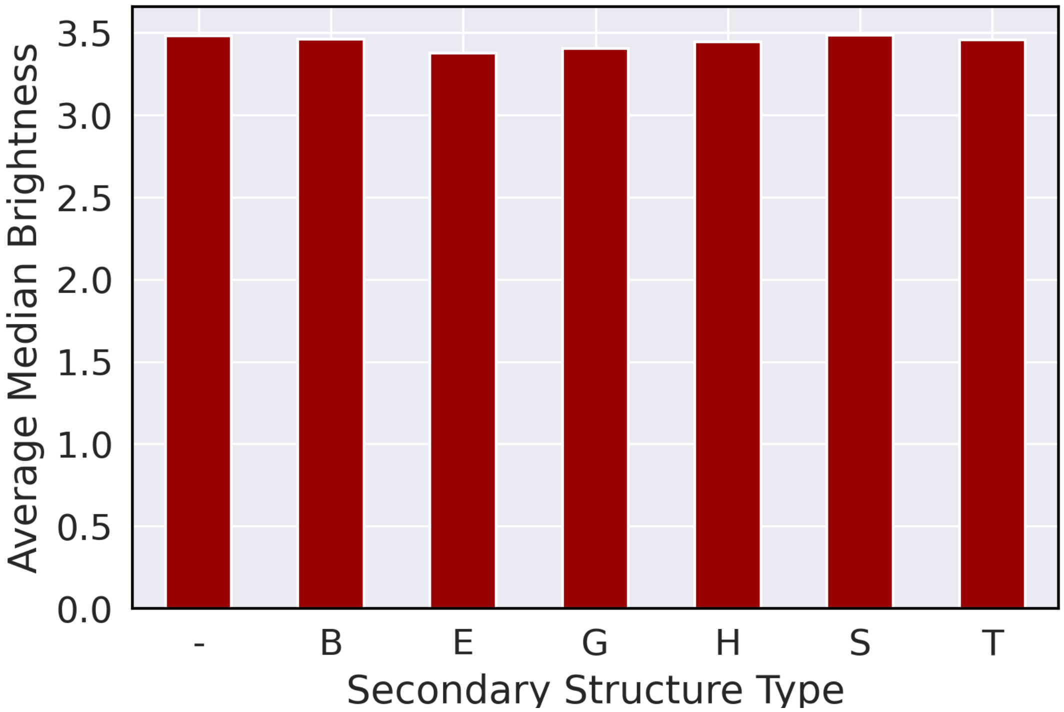


**Figure S6.** Bar chart of the average brightness per secondary structure type

**Table S4.** Table displaying statistics of secondary structure on the Sarkisyan dataset.

| Secondary Structure | Minimum | Maximum | Mean | Stdev |
|---|---|---|---|---|
| - | 1.301 | 3.907 | 3.479 | 0.568 |
| B | 1.401 | 3.800 | 3.460 | 0.606 |
| E | 1.301 | 4.114 | 3.377 | 0.691 |
| G | 1.299 | 3.992 | 3.406 | 0.736 |
| H | 1.528 | 3.766 | 3.447 | 0.548 |

| | | | | |
|---|---|---|---|---|
| S | 1.301 | 3.798 | 3.485 | 0.478 |
| T | 1.301 | 3.804 | 3.456 | 0.537 |

The RecA dataset was analyzed for discovering the associations between the mutation type and the corresponding phenotype change. We wrote a Python script to calculate the percentage of each phenotype as a function of mutated amino acid type. Figure S7 is a bar chart for each unique amino acid mutation type in the dataset.

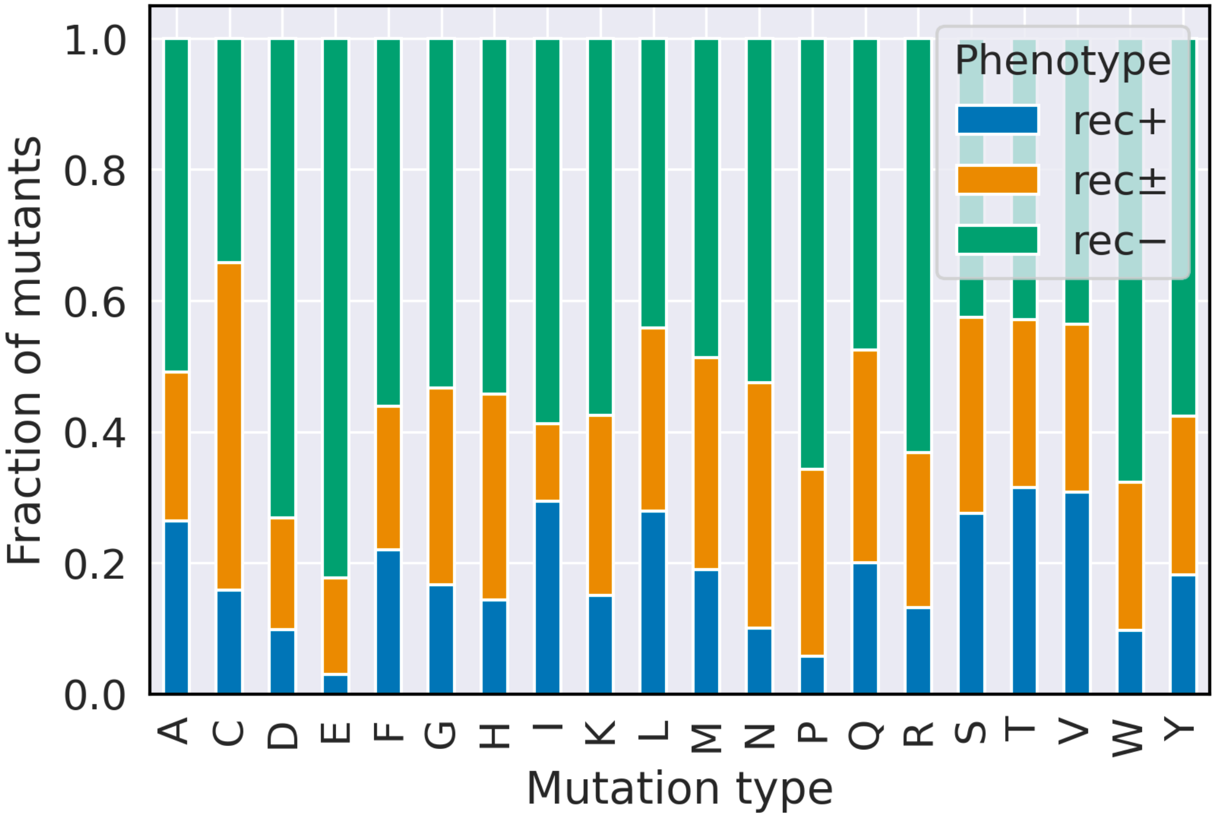


**Figure S7**. Bar chart the percentage of phenotype per mutation type in the RecA dataset.

This percentage bar plot shows that mutation to glutamic acid most likely leads to misfunction of RecA proteins, which possibly inhibits the SOS response by deactivating RecA and then preventing LexA from self-cleavage. In addition to glutamic acid, aspartic acid also has a significant effect on disabling the functionality of RecA protein. We provide statistics about the percentages of phenotype per mutation type in Table S5.

**Table S5.** Table displaying statistics of the resultant percentages of phenotype per mutation type on the RecA dataset.

| Mutation Type | RecA- 2 | RecA± 1 | RecA+ 0 |
|---|---|---|---|
| A | 50.94% | 22.64% | 26.42% |
| C | 34.21% | 50.00% | 15.79% |
| D | 73.17% | 17.07% | 9.756% |
| E | 82.35% | 14.71% | 2.94% |
| F | 56.10% | 21.95% | 21.95% |
| G | 53.33% | 30.00% | 16.67% |
| H | 54.29% | 31.43% | 14.28% |
| I | 58.82% | 11.77% | 29.41% |
| K | 57.50% | 27.50% | 15.00% |
| L | 44.19% | 27.91% | 27.90% |
| M | 48.65% | 32.43% | 18.92% |
| N | 52.50% | 37.50% | 10.00% |
| P | 65.71% | 28.57% | 5.72% |
| Q | 47.50% | 32.50% | 20.00% |

| | | | |
|---|---|---|---|
| R | 63.16% | 23.68% | 13.16% |
| S | 42.50% | 30.00% | 27.50% |
| T | 42.86% | 25.71% | 31.43% |
| V | 43.59% | 25.64% | 30.77% |
| W | 67.74% | 22.58% | 9.68% |
| Y | 57.58% | 24.24% | 18.18% |

The same analyses were performed for secondary structure motifs. Figure S8 displays a bar chart of the resultant percentages of phenotype per secondary structure type. In addition, we provide statistics of phenotype percentages for secondary structure motifs in Table S6.

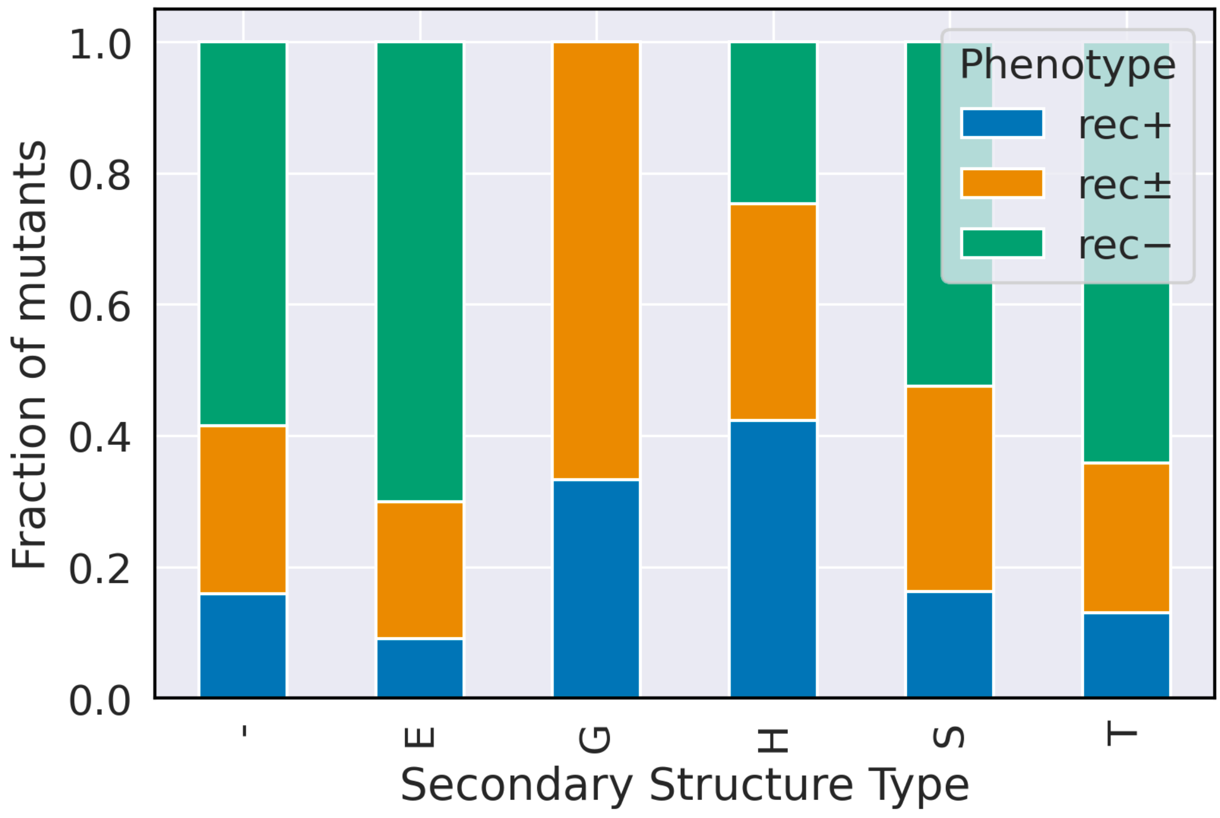

**Figure S8.** Bar chart plotting of the resultant percentages of phenotype per secondary structure type.

**Table S6.** Table displaying statistics of the resultant percentages of phenotype per mutation type on the RecA dataset.

| Secondary Structure | RecA- | RecA± | RecA+ |
|---|---|---|---|
| - | 58.52% | 25.57% | 15.91% |
| E | 70.05% | 20.86% | 9.09% |
| G | 0 | 66.67% | 33.34% |
| H | 24.62% | 33.08% | 42.30% |
| S | 52.41% | 31.32% | 16.27% |
| T | 64.13% | 22.83% | 13.04% |

**Hint token investigation**

To further investigate the effect of the hint tokens on PLMs beyond attention plot analysis, we visualized model embeddings. The embeddings for both the ProtHTL_ΔG and Prot_ΔG models were reduced to two dimensions using the UMAP implementation from CuML. Figure S9 and Figure S10 display these model embeddings colored by energy bin labels and the number of mutations in the sequence.

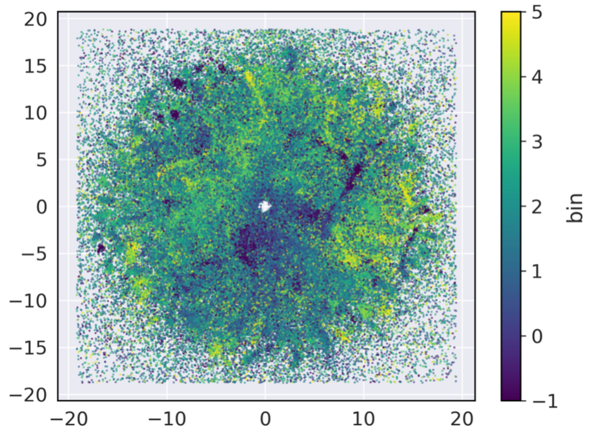


**Figure S9.** UMAP embeddings for ProtHTL_ΔG model colored by experimental label bins and by number of mutations.

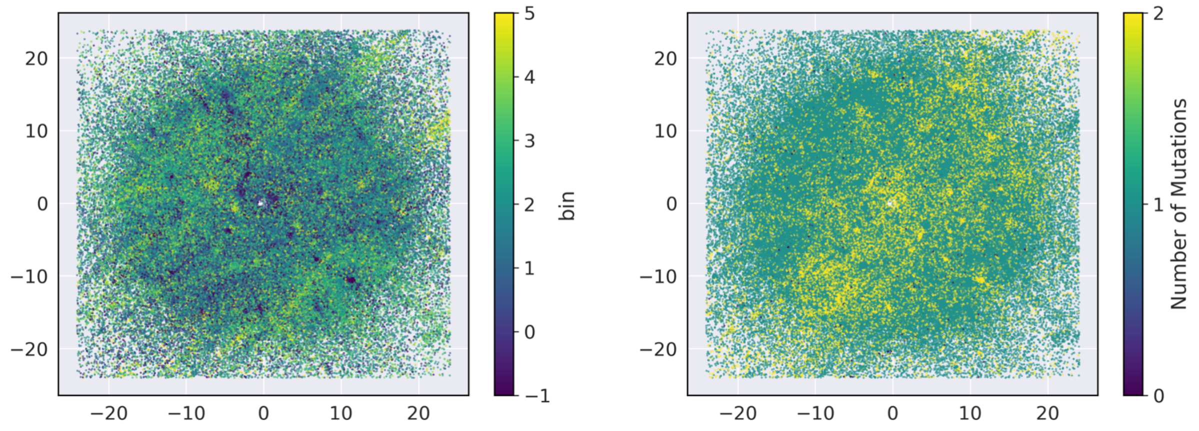


**Figure S10.** UMAP embeddings for Prot_ΔG model colored by experimental label bin and by number of mutations.

Here, we observed that datapoints with different ΔG energy ranges are distributed over the embedding space. Also, we observed that WT, single, and double mutations could be found across the entirety of the embedding space. However, when comparing the embeddings between the HTL and traditional finetuning model, we find that these embeddings are of a totally different shape.

This simply indicates that they see the data differently, which is supported by the attention plot analysis.

We also confirmed the similar trend in Sarkisyan dataset and RecA dataset in Figure S11-18 that models with hint token application see data differently from models without hint token application. In addition, models with the application of hint tokens have better clustering of datapoints with same labels compared with models without the application of hint tokens.

**Figure S11**. UMAP embeddings for Prot_ΔG_GFP model, Prot_GFP model, ProtHTL_ΔG_GFP model, and ProtHTL_GFP model trained on 1% Sarkisyan dataset, colored by experimental label bins.

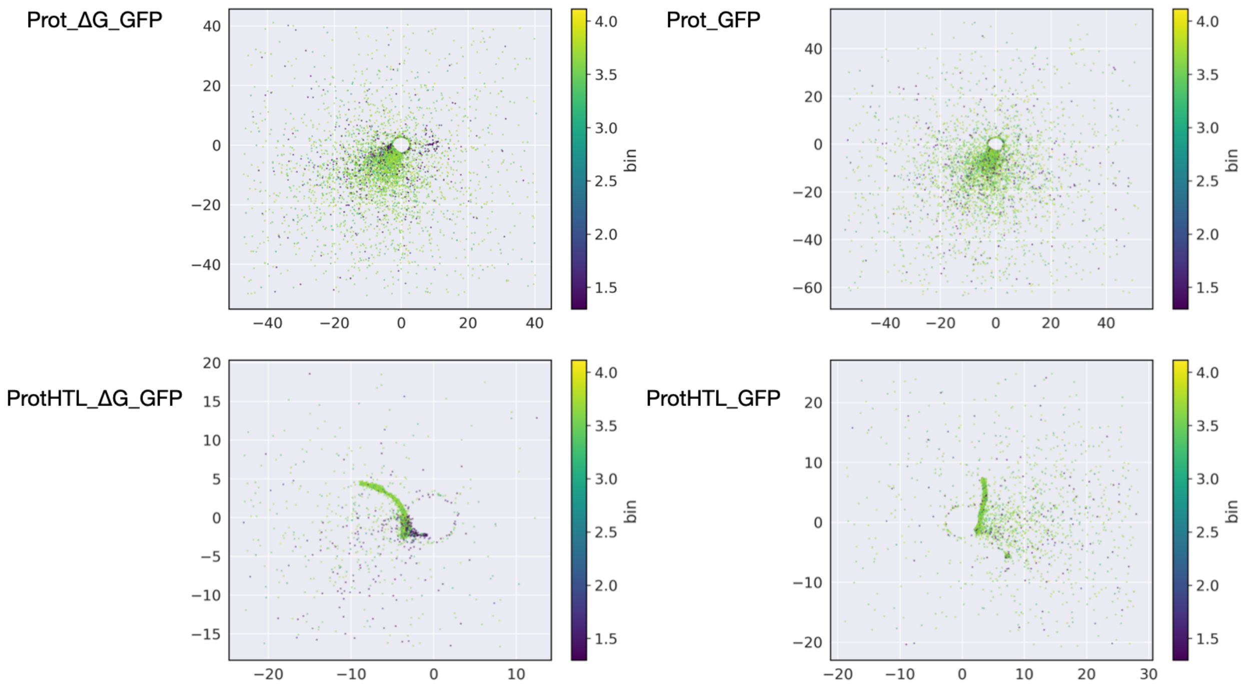


**Figure S12.** UMAP embeddings for Prot_ΔG_GFP model, Prot_GFP model, ProtHTL_ΔG_GFP model, and ProtHTL_GFP model trained on 1% Sarkisyan dataset, colored by the number of mutations.

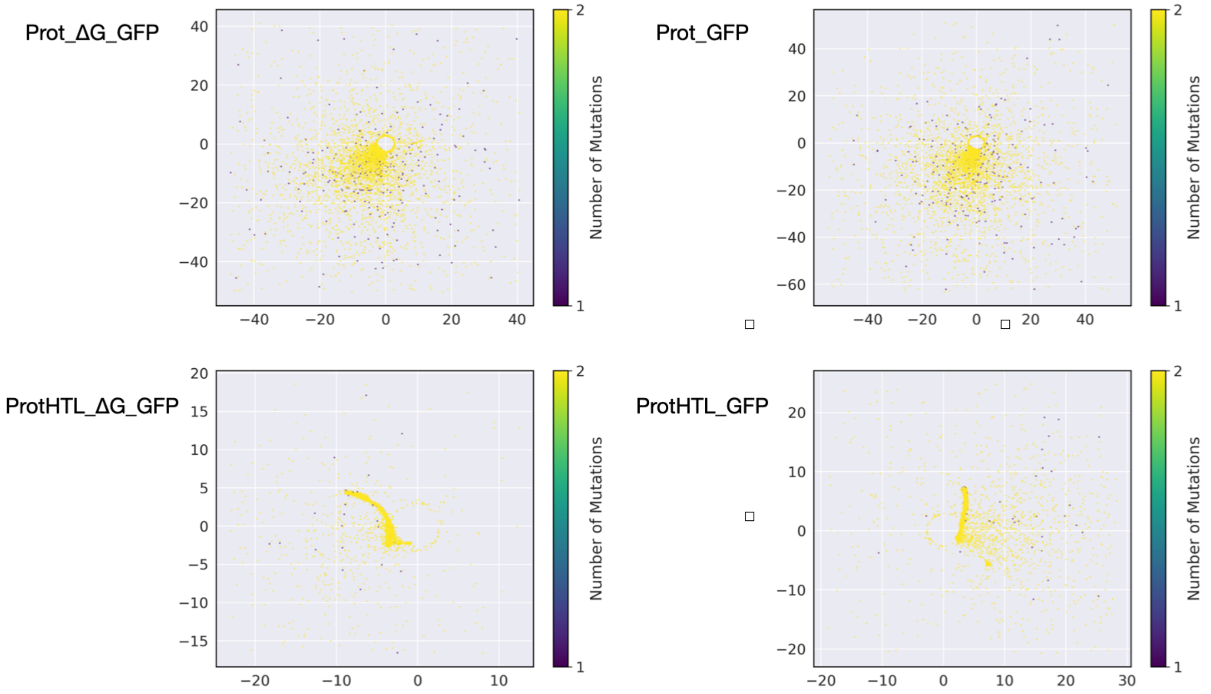


**Figure S13.** UMAP embeddings for Prot_ΔG_GFP model, Prot_GFP model, ProtHTL_ΔG_GFP model, and ProtHTL_GFP model trained on 10% Sarkisyan dataset, colored by experimental label bins.

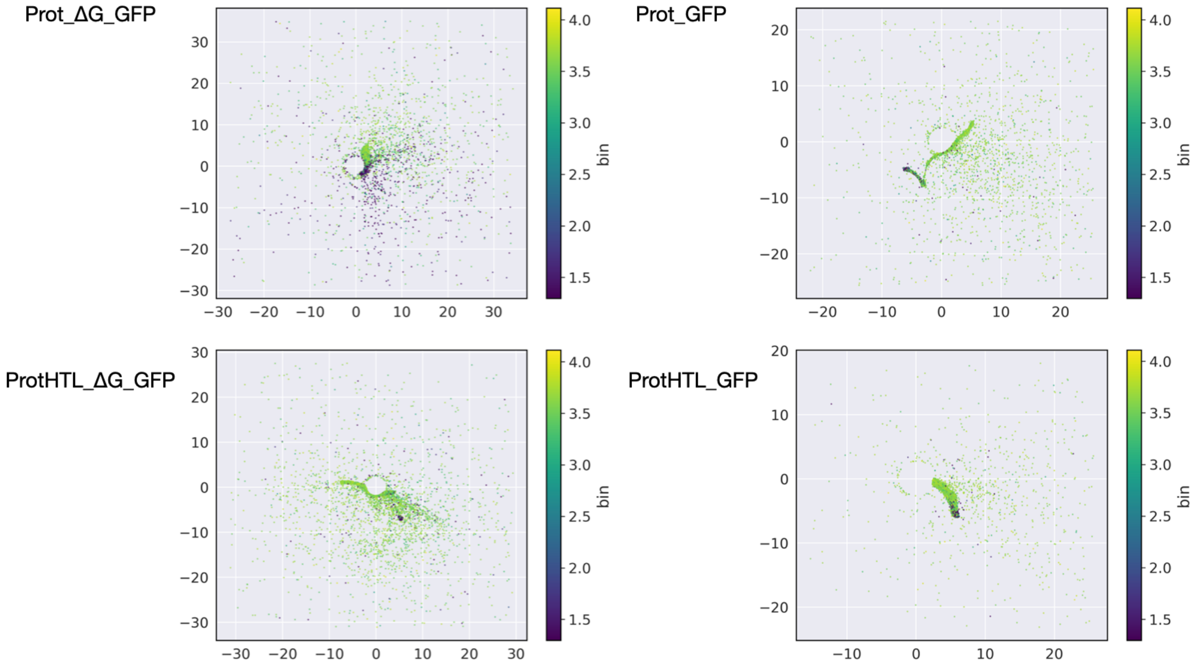


**Figure S14.** UMAP embeddings for Prot_ΔG_GFP model, Prot_GFP model, ProtHTL_ΔG_GFP model, and ProtHTL_GFP model trained on 10% Sarkisyan dataset, colored by the number of mutations.

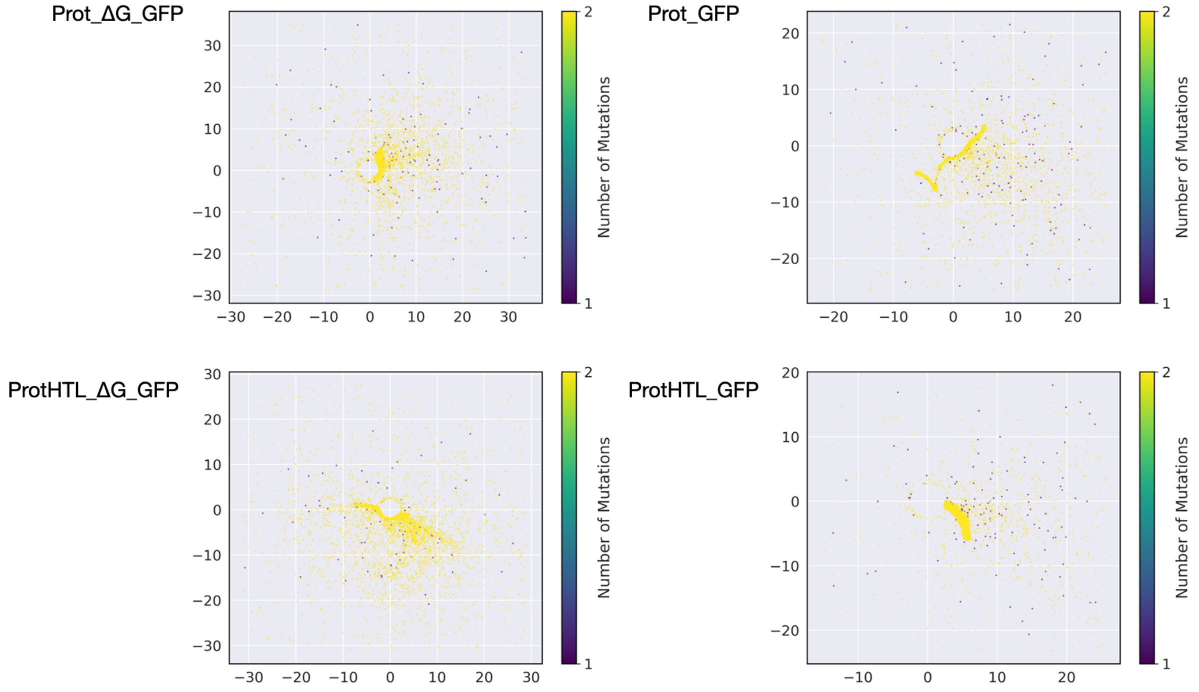


**Figure S15.** UMAP embeddings for Prot_ΔG_GFP model, Prot_GFP model, ProtHTL_ΔG_GFP model, and ProtHTL_GFP model trained on 80% Sarkisyan dataset, colored by experimental label bins.

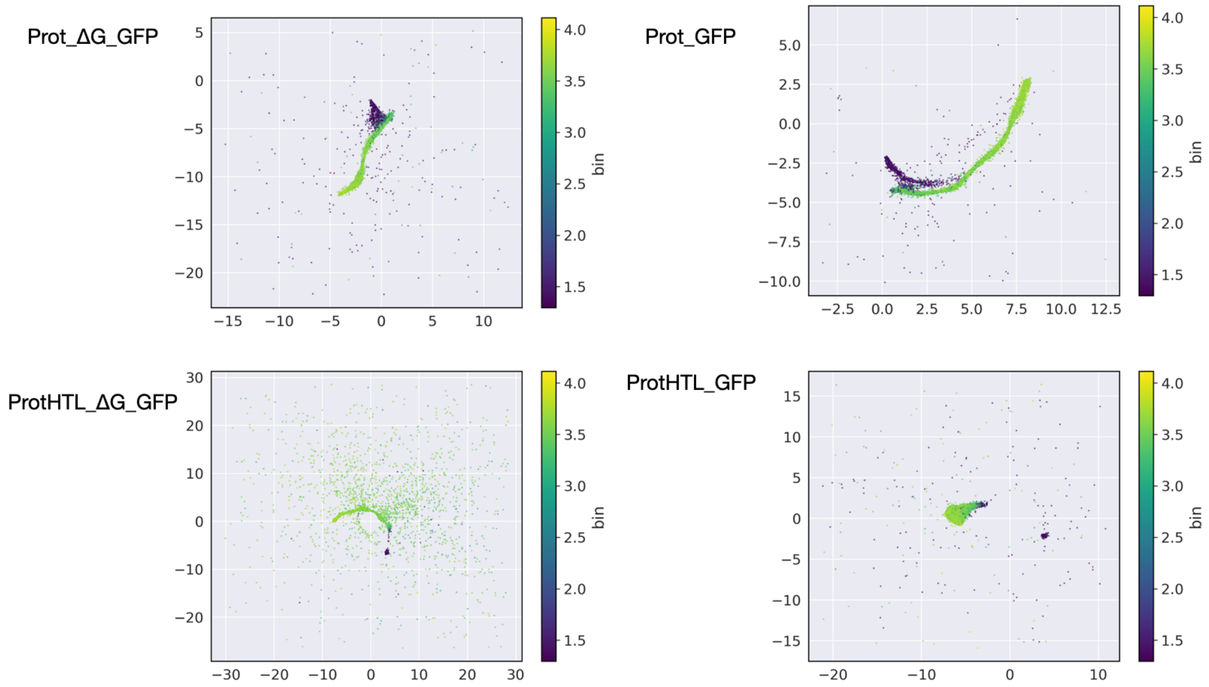


**Figure S16.** UMAP embeddings for Prot_ΔG_GFP model, Prot_GFP model, ProtHTL_ΔG_GFP model, and ProtHTL_GFP model trained on 80% Sarkisyan dataset, colored by the number of mutations.

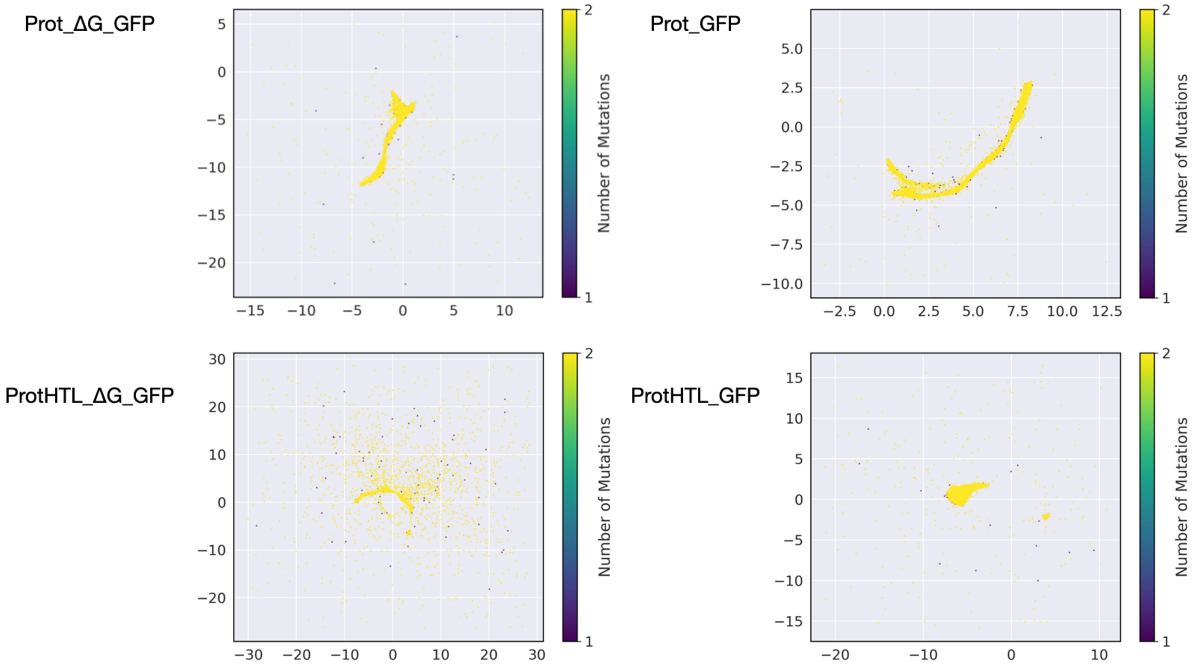


**Figure S17.** UMAP embeddings for Prot_ΔG_RecA model, Prot_RecA model, ProtHTL_ΔG_RecA model, and ProtHTL_RecA model, colored by phenotype labels.

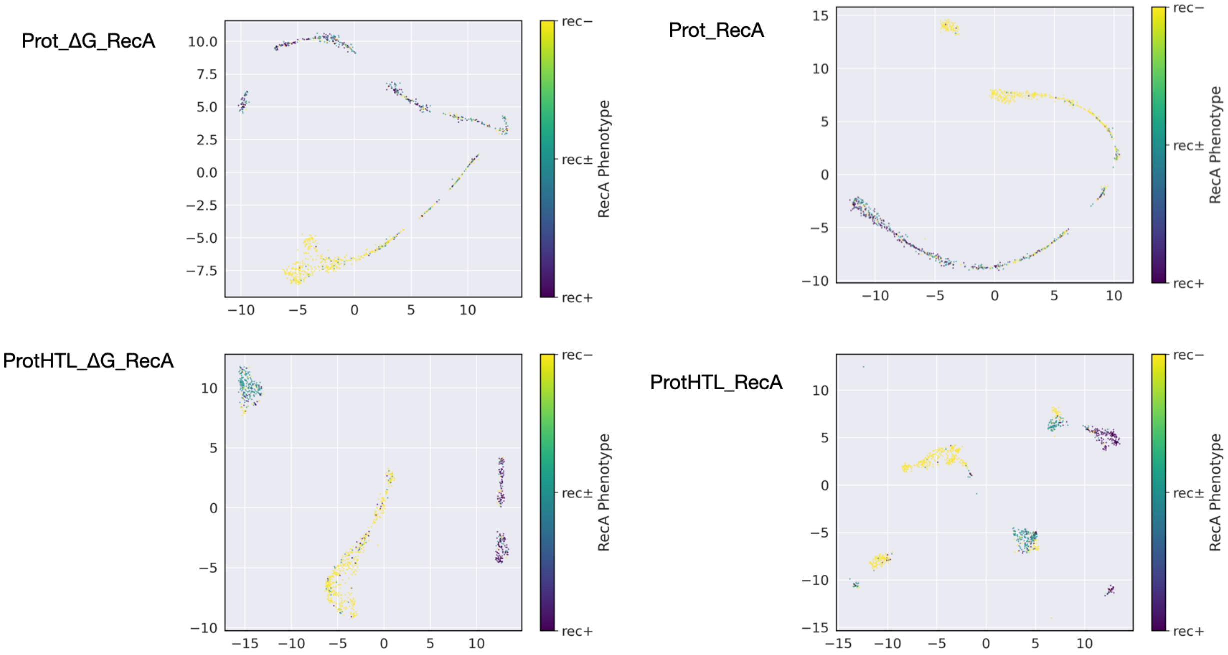

**Figure S18.** UMAP embeddings for Prot_ΔG_RecA model, Prot_RecA model, ProtHTL_ΔG_RecA model, and ProtHTL_RecA model, colored by the number of mutations.

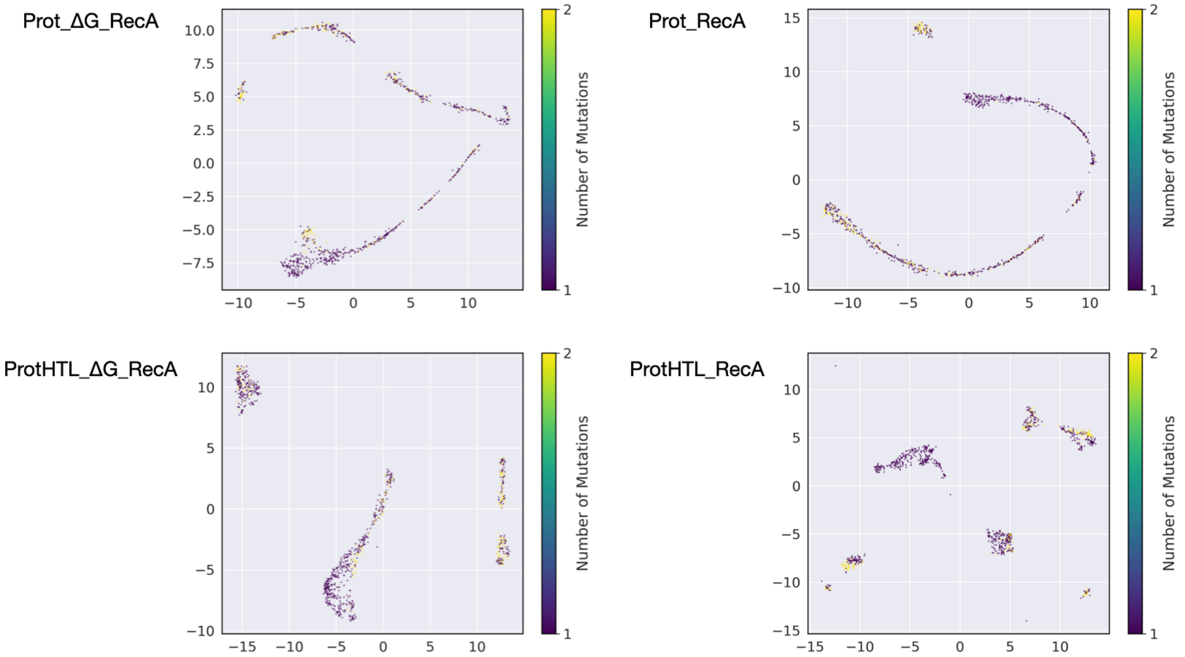

### Model Training

Hyperparameter Tuning

Our models are hyperparameter-tuned by Optuna Bayesian optimization and the best parameters for all the models are shown in Table S7.

**Table S7.** Best hyperparameters for all models obtained by Optuna Bayesian optimization

| | weight_decay | batch_size | learning_rate | random_seed |
|---|---|---|---|---|
| ProtHTL_ΔG | 0.004 | 128 | 8.369042894376066e-05 | 44 |
| Prot_ΔG | 0.006 | 128 | 6.746417134006616e-06 | 42 |
| ProtHTL_ΔG_GFP(1%) | 0.005 | 32 | 3.6934741669941536e-05 | 40 |
| ProtHTL_GFP(1%) | 0.01 | 32 | 6.923767077310682e-06 | 40 |
| Prot_ΔG_GFP(1%) | 0.007 | 32 | 6.832986140924483e-06 | 40 |
| Prot_GFP(1%) | 0.01 | 32 | 0.00047840373722658096 | 42 |

| | | | | |
|---|---|---|---|---|
| ProtHTL_ΔG_GFP(10%) | 0.007 | 32 | 2.814914808453654e-05 | 42 |
| ProtHTL_GFP(10%) | 0.01 | 32 | 5.351732463689635e-06 | 42 |
| Prot_ΔG_GFP(10%) | 0.01 | 32 | 7.98456596918626e-06 | 40 |
| Prot_GFP(10%) | 0.004 | 32 | 9.89394320267359e-06 | 42 |
| ProtHTL_ΔG_GFP(80%) | 0.009000000000000001 | 64 | 3.396426173109566e-05 | 40 |
| ProtHTL_GFP(80%) | 0.003 | 64 | 5.176838813579891e-06 | 40 |
| Prot_ΔG_GFP(80%) | 0.008 | 64 | 1.6895054770217645e-05 | 40 |
| Prot_GFP(80%) | 0.002 | 64 | 1.3472091288824858e-05 | 40 |
| ProtHTL_ΔG_RecA | 0.001498383300575274 | 14 | 1.0724513869451559e-05 | 42 |
| ProtHTL_RecA | 0.0012725552529390396 | 12 | 1.6587909547103302e-05 | 42 |

| | | | | |
|---|---|---|---|---|
| Prot_ΔG_RecA | 0.0007501418504121116 | 10 | 1.1049769146353426e-05 | 42 |
| Prot_GFP_RecA | 0.0010199291067622914 | 12 | 1.0327209030070408e-05 | 42 |

Metrics for validation sets

Table S8- 10 are validation metrics for Tsuboyama dataset, Sarkisyan dataset, and RecA dataset.

**Table S8.** Validation metrics Tsuboyama dataset.

| | Prot_ΔG | ProtHTL_ΔG |
|---|---|---|
| Finetuning method | Traditional | HTL |
| $R^2$ | 0.37 | 0.45 |

**Table S9.** Validation metrics Sarkisyan dataset.

| | Prot_GFP | ProtHTL_GFP | Prot_ΔG_GFP | ProtHTL_ΔG_GFP |
|---|---|---|---|---|
| Pretrained model | ProtBERT | ProtBERT | Prot_ΔG | ProtHTL_ΔG |
| Finetuning method | Traditional | HTL | Traditional | HTL |
| Training Data (%) | ($R^2$) | ($R^2$) | ($R^2$) | ($R^2$) |
| 1 | NaN | 0.36 | 0.37 | 0.71 |
| 10 | 0.32 | 0.28 | 0.72 | 0.71 |

| | | | | |
|---|---|---|---|---|
| 80 | 0.83 | 0.87 | 0.88 | 0.86 |

**Table S10.** Validation metrics RecA dataset.

| | Prot_RecA | ProtHTL_RecA | Prot_ΔG_RecA | ProtHTL_ΔG_RecA |
|---|---|---|---|---|
| Pretrained model | ProtBERT | ProtBERT | Prot_ΔG | ProtHTL_ΔG |
| Finetuning method | Traditional | HTL | Traditional | HTL |
| Accuracy | 0.62 | 0.73 | 0.64 | 0.68 |
| Precision | 0.66 | 0.77 | 0.69 | 0.74 |
| Recall | 0.68 | 0.76 | 0.69 | 0.74 |
| F1 | 0.65 | 0.76 | 0.68 | 0.74 |
